# Macrophages redeploy functional cancer cell surface proteins following phagocytosis

**DOI:** 10.1101/2024.09.23.613776

**Authors:** Regan F. Volk, Sara W. Casebeer, Andrew C. Condon, Bahar Zirak, Nayelis Manon, Iryna Irkliyenko, Huajun Liao, Shao Tao, Tommaso Pollini, Vijay Ramani, Ajay V. Maker, Trevor Fidler, Hani Goodarzi, Balyn W. Zaro

**Affiliations:** Department of Pharmaceutical Chemistry, The Cardiovascular Research Institute, Helen Diller Family Comprehensive Cancer Center, Quantitative Biosciences Institute, School of Pharmacy, University of California, San Francisco, CA, USA; Department of Biochemistry and Biophysics, Department of Urology, Helen Diller Comprehensive Cancer Center, Bakar Computational Health Science Institute, University of California, San Francisco, CA, USA; Arc Institute, Palo Alto, CA, USA; Gladstone Institute for Data Science and Biotechnology, San Francisco, CA, USA; Department of Biochemistry and Biophysics, Helen Diller Comprehensive Cancer Center, Bakar Computational Health Science Institute, University of California, San Francisco, CA, USA; The Cardiovascular Research Institute, University of California, San Francisco, CA, USA; Division of Surgical Oncology, Department of Surgery, University of California, San Francisco, CA, USA

## Abstract

Macrophage-mediated phagocytosis is a vital innate immune process altered in cancer. We show here that tumor-associated macrophages (TAMs) redeploy intact cell surface proteins from cancer cells to their own cell surface. We initially observed the canonical epithelial cancer surface marker EpCAM on the surface of TAMs in primary human solid tumors but not paired peripheral blood macrophages. In a murine model of metastatic breast cancer, we also observed EpCAM on the surface of primary TAMs that have phagocytosed breast cancer cells. In a model of a myeloproliferative neoplasm, we again found engulfed cell-derived surface proteins on the surface of macrophages following phagocytosis. A co-culture system and proteomics assay that tags proteins based on their cell-of-origin revealed hundreds of cell surface proteins synthesized in cancer cells are present and fully intact on the surface of macrophages following phagocytosis. Using a biotin transfer assay, we determined that these proteins were on the surface of the cancer cell prior to redeployment by the macrophage following phagocytosis. Furthermore, macrophages that redeploy a neutral amino acid transporter correspondingly show increased transport of an unnatural amino acid substrate. Widespread acquisition of proteins from engulfed cells may contribute to two critical TAM phenotypes: the inability to phagocytose and reprogrammed metabolism.

## Main Text

Despite contributing significantly to solid tumor bulk and being associated with poor disease outcomes, tumor associated macrophages (TAMs) have yet to be targeted by a clinically-approved therapeutic^1,2^. TAMs comprise a diverse cell phenotype and are often problematic in the tumor microenvironment, suppressing immune cell activation, increasing tumor density, promoting tumor growth and metastasis, and supporting angiogenesis^3^. Like cancer cells, at least a subset of TAMs undergo a metabolic shift in the acidic tumor microenvironment, further driving their dysregulation^4–8^. Metabolic crosstalk between TAMs and cancer cells has also been reported^9^. Moreover, TAMs are poor phagocytes, unlike recently recruited peripheral blood macrophages^10^. Much effort has been expended to reprogram TAMs or deplete them from tumors as potential innate immunotherapeutics^11,12^. However, an understanding of the etiology of the TAM remains elusive, hampering development of TAM-targeting therapeutics.

Macrophages are capable of trogocytosing neighboring cell surface proteins leading to the incorporation of proteins on their surface that were synthesized by a nearby cell^13,14^. Macrophages can also acquire cell surface proteins from nearby cells through an Fc-receptor-antibody mediated shaving mechanism^15,16^. RNA-sequencing from solid tumors often reveals transcriptional signatures for a subset of TAMs akin to neighboring cancer cells^5,6,17–19^. It remains unclear if this RNA signature results in a novel subset of proteins synthesized in macrophages (where transcript was synthesized in either the cancer cell or macrophage), or if these transcripts correspond to residual surviving RNA following phagocytosis of a cancer cell. Beyond revealing a fundamental feature of cell-intrinsic TAM biology, validating if and how intact cancer cell proteins appear on the surface of TAMs could also reshape our understanding of the tumor microenvironment and clarify the role of TAMs in activating or suppressing antigen-specific T-cells. Therefore, we set out to investigate if TAMs were expressing proteins normally synthesized in cancer cells and, if so, how this occurs.

Single-cell suspensions from Epithelial cell adhesion molecule (EpCAM)-positive solid tumor resections revealed TAMs staining positive for EpCAM (Figure 1A, Extended Data 1). Paired peripheral blood macrophages from these patients did not stain positive for EpCAM and neither did monocyte-derived macrophages (MDM), suggesting a mechanism specific to the tumor microenvironment (Figure 1A, Extended Data Figure 1). Having observed this established pro-growth epithelial cancer cell marker on the surface of primary human TAMs^20,21^, we next asked if EpCAM could be observed in the established transplantable and metastatic 4T1 mouse mammary carcinoma^22,23^. Using FACS, we observed a subset of EpCAM^+^ TAMs, and purified EpCAM^+^ and EpCAM^−^ TAMs from primary tumors (Figure 1B, Extended Data Figure 2). By mass spectrometry (MS)-based proteomic analysis, we showed that EpCAM is exclusively detected in EpCAM^+^ samples, confirming the purity of our EpCAM-dependent sort (Table 1). We next compared the relative abundance of proteins detected in EpCAM^+^ TAMs and EpCAM^−^ TAMs. We cross-referenced these proteins to the whole proteomes of EpCAM^+^ cancer cells also isolated and purified from these tumors. A statistically significant enrichment of 4T1 cancer cell proteins was detectable in EpCAM^+^ TAMs compared to EpCAM^−^ TAMs (Figure 1C). Annotation of the subcellular localization of proteins detected in EpCAM^+^ TAMs likely to have come from 4T1 cells revealed myriad intracellular proteins from cell compartments including but not limited to the nucleus, cytosol, and mitochondria (Extended Data Figure 3A). Proteins attributed to 4T1 cells were not heavily biased towards the most abundant 4T1 proteins (Median Percentile 53%, Extended Data Figure 3B). These MS data implicate whole cell uptake (i.e. phagocytosis) rather than trogocytosis of cancer cell membranes by TAMs ^15,16^. Publicly available scRNA-Seq data indicates that EpCAM RNA is detectable in a subset of macrophages, perhaps indicating these cells have recently phagocytosed cancer cells (Extended Data Figure 4) ^24^. Taken together, our data show that macrophages that have phagocytosed target cells express a cell surface protein uniquely encoded by the phagocytosed cell, namely EpCAM.

**Figure 1.**
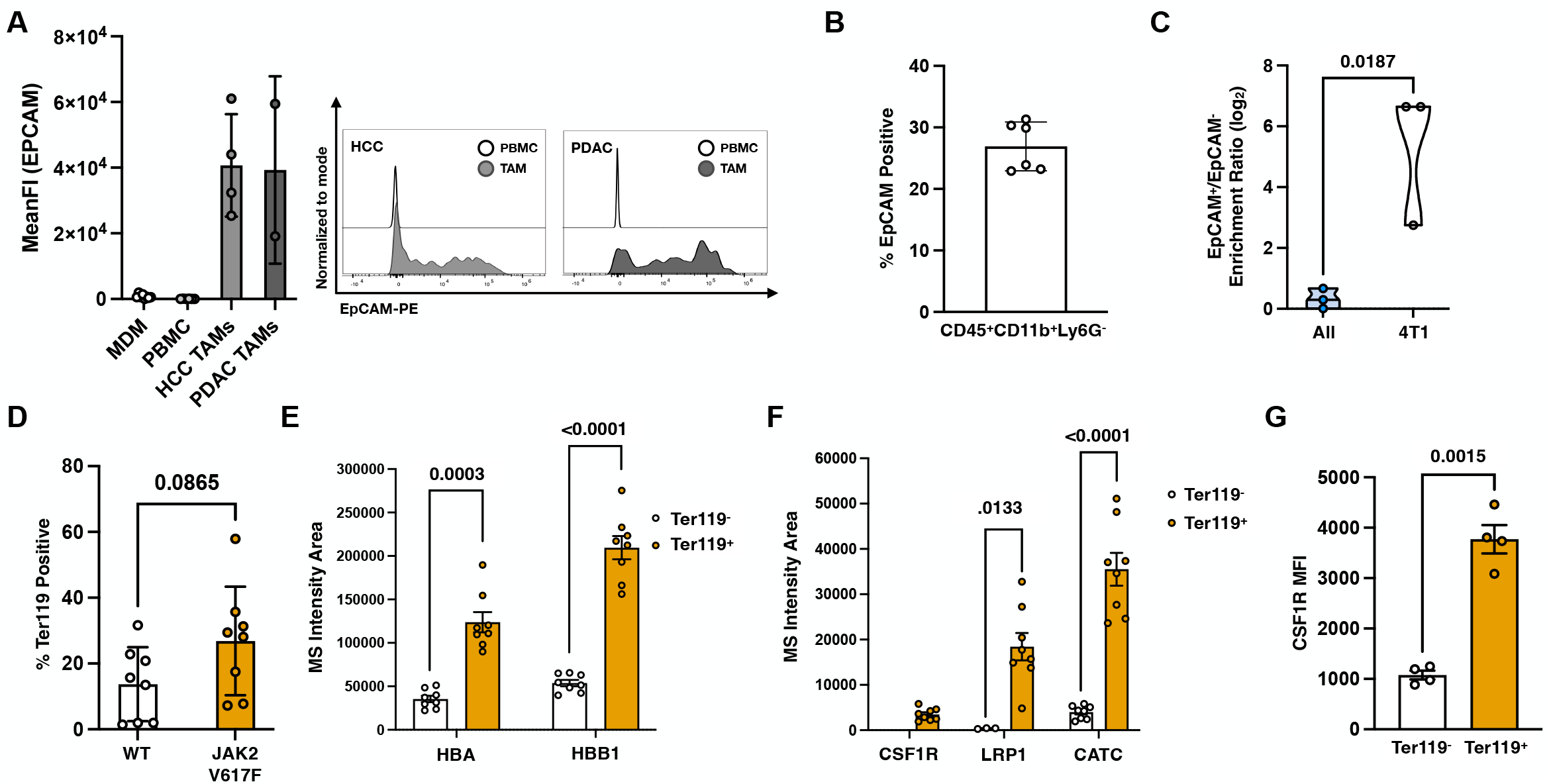
Established surface markers of cancer cells and red blood cells are detectable on the surface of primary macrophages that have phagocytosed. A. Anti-EpCAM staining of tumor associated macrophages (TAMs, CD45^+^ CD11b^+^ CD14^+^) from human hepatocellular carcinoma (HCC) or pancreatic ductal adenocarcinoma (PDAC) resections, matched peripheral blood macrophages (PBMC), or monocyte derived macrophages (MDM). n = 3 biological replicates (PMBC), 4 biological replicates (MDM), 2 biological replicates (HCC), 1 biological replicate (PDAC), 2 technical replicates per biological replicate. Representative histograms for EpCAM staining from HCC or PDAC TAMs and paired PBMC macrophages. B. Frequency of EpCAM positive macrophages (CD45^+^ CD11b^+^ Ly6G^-^) isolated from dissociated 4T1 tumors. n = 6 biological replicates. C. Median intensity ratio for either all proteins detectable in CD45^+^ CD11b^+^ Ly6G^-^EpCAM^+^ vs CD45^+^ CD11b^+^ Ly6G^-^EpCAM^-^ cells or subset of proteins determined to be of non-macrophage and likely 4T1 cancer cell origin. n = 3 replicates D. Ter119 positivity of spleen-resident macrophages as determined by flow cytometry in wild-type and JAK2 V617F mutant mice. n = 4 biological, 2 technical replicates each. E. Mass spectrometry detection of the red blood cell proteins HBA and HBB1 detectable in Ter119^+^ and Ter119^-^ macrophages from JAK2 V617F mice. n = 4 biological replicates, 2 technical replicates each. F. Mass spectrometry detection of macrophage proteins CSF1R, LRP1, and CATC in Ter119^+^ and Ter119^-^ macrophages from JAK2 V617F mice. n = 4 biological replicates, 2 technical replicates. G. CSF1R mean fluorescence intensity (MFI) of macrophages in the spleens of JAK2 V617F mice n = 5 biological replicates. Error bars denote standard deviation.

We tested if macrophages in a mouse model of a blood pre-malignancy also have proteins on their cell surface characteristic of their phagocytic target. We selected the myeloproliferative neoplasm JAK2 V617F model because this mutant displays increased macrophage-mediated phagocytosis of red blood cells (RBCs) in the spleen^25^. We tested macrophages from the spleens of JAK2 V617F mice for Ter119, an RBC-specific surface epitope (Figure 1D, Extended Data Figure 5). Notably, the reported increase in phagocytosis observed between wild-type mice and the JAK2 V617F mutant was consistent with the increase in relative Ter119 positivity in the macrophage population^26^. Ter119^+^ and Ter119^-^ macrophages were purified by FACS and subjected to MS analysis, revealing that Ter119^+^ macrophages had higher levels of RBC proteins in their whole cell proteomes (Tables 2 and 3)^26^. Established intracellular RBC proteins HBA and HBB1 were more detectable in Ter119^+^ macrophages compared to Ter119^-^ macrophages (Figure 1E). This observation suggested again that surface proteins from cells phagocytosed by macrophages are detectable on the macrophage cell surface. Thus, as with the solid tumor paradigm, the JAK2 V617F macrophages display a surface epitope characteristic of the target cell in a manner that correlates with prior phagocytosis of the target cell, as inferred from the presence of abundant target cell proteins.

We also observed surprising differences in the detection of proteins that are known to be encoded and synthesized by macrophages, suggesting that there may be functional and/or cell-state differences between Ter119^+^ and Ter119^-^ macrophages in the spleen (Figure 1F, Tables 2 and 3). Specifically, CSF1R and CATC expression have previously been implicated in phagocytic capacity and macrophage polarization^27,28^, and LRP1 is an established receptor mediating macrophage phagocytosis of live or apoptotic red blood cells^29^. We further validated differential CSF1R abundance between Ter119^+^ and Ter119^-^ macrophages by flow cytometry (Figure 1G, Extended Data Figure 6).

To rigorously investigate the possibility that surface proteins are transferred to the macrophage from the target cell as a consequence of phagocytosis, we devised a short 2-hour co-culture assay coupled to an MS method that enables assignment of each protein’s cell of origin. Primary human monocyte-derived macrophages were challenged to phagocytose the human colorectal cancer cell line SW620 stained with an intracellular viability dye. This allowed us to purify macrophages that had phagocytosed live cancer cells (“Eaters”) and those which had not (“Non-Eaters”) by FACS (Figure 2A, Extended Data Figure 7)^30,31^. To distinguish macrophage and cancer cell protein contributions from Eater macrophages and validate the purity of our samples, we repurposed isotopic chemical tags to assign each protein’s origin^32^. Proteins synthesized in the cancer cell were isotopically heavier than those synthesized in the macrophage. To confirm that our cell populations were pure beyond flow cytometry plots, we subjected each population to RNA-sequencing and MS analysis.

**Figure 2.**
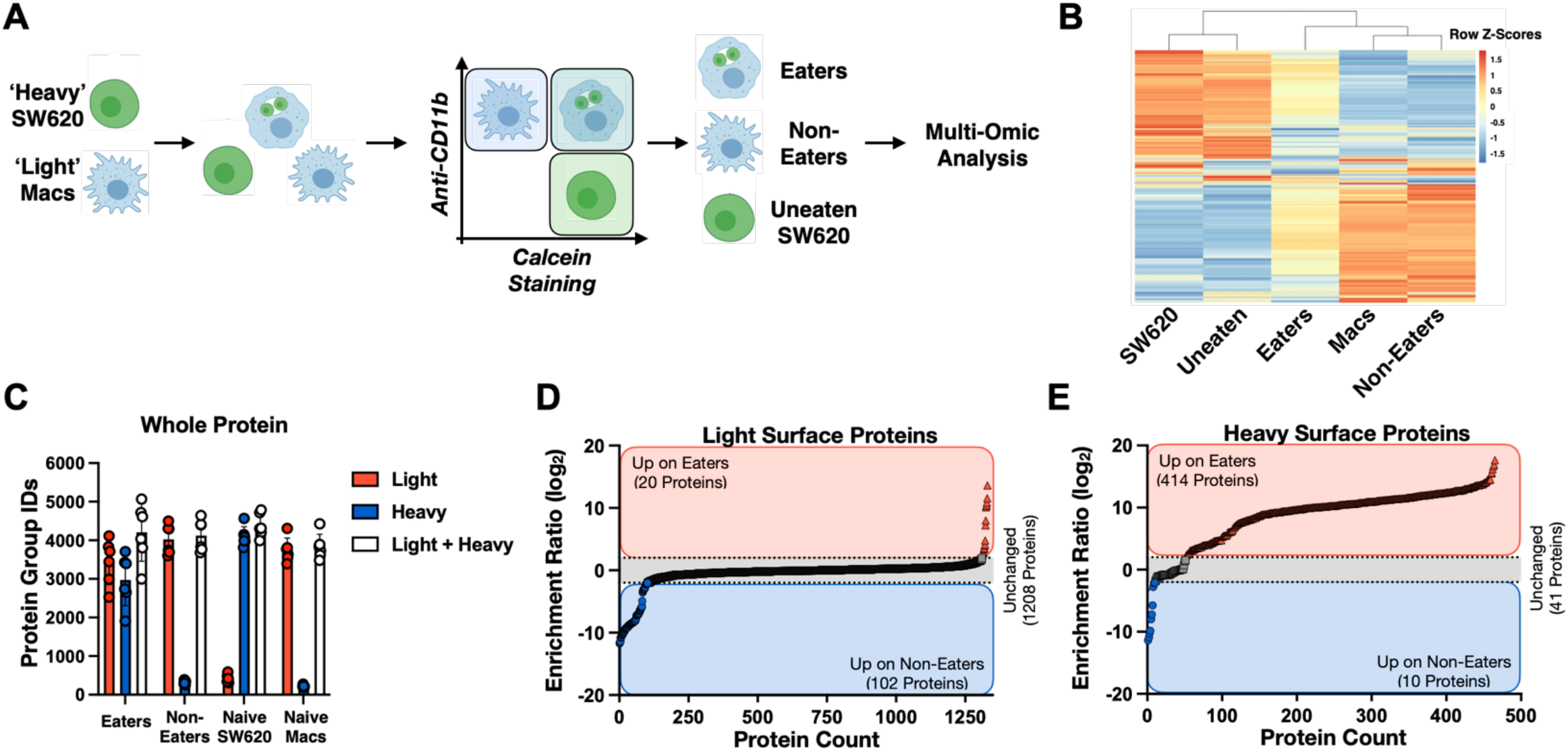
Mass spectrometry analysis of a macrophage co-culture system reveals that macrophages that phagocytose colorectal cancer cells incorporate cancer cell-surface proteins onto their own cell surface. A. Experimental workflow. Donor-derived macrophages are incubated with isotopically heavy, calcein stained SW620 colorectal cancer cells (2 h, 37°C). Cell populations (Eaters, Non-Eaters, and Uneaten) are purified via FACS using the macrophage marker CD11b and calcein. Sorted cells are prepared for mass spectrometry (MS) analysis or RNA-sequencing. For surface proteomics, cell populations are labeled with a positively-charged biotin reagent prior to enrichment. B. Heat map of DESeq2 normalized log-transformed gene counts across naive cells or those purified following co-incubation. Z-scores generated by scaling across rows. n = 2 biological replicates, 2 technical replicates. C. Total proteins identified when performing MS search for light-only proteins (no modification), heavy-only proteins (fixed modification), or light and heavy (variable modification). Error bars are shown as SD. D. Relative abundance of light surface proteins detectable on Eaters compared to Non-Eaters. E. Relative abundance of heavy surface proteins detectable on Eaters compared to Non-Eaters.

Naive macrophages unchallenged with cancer cells and Non-Eater macrophages in co-culture were transcriptionally similar (Figure 2B, Table 4). Naive SW620 cancer cells and those in co-culture that were not phagocytosed (Uneaten), were also similar to each other (Figure 2B, Table 4). Conversely, Eater macrophages appeared as transcriptionally distinct hybrids of macrophages and cancer cells, validating that mRNA from cancer cells may remain viable for some period of time following phagocytosis. MS analysis further confirmed the purity of each cell population. Eater macrophages harbored both heavy and light proteins, indicating the presence of proteins synthesized in both macrophages and cancer cells, while Non-Eater macrophages contained mostly light proteins (Figure 2C, Tables 5A-C). SW620 cancer cells contained mostly heavy proteins.

Having confirmed the purity of Eaters and Non-Eaters as macrophages that have phagocytosed or not phagocytosed live cancer cells, respectively, we further optimized our assay to be able to purify and comparatively characterize the cell surface proteome. A vast majority of the over 1200 light proteins enriched from the surface of Eaters and Non-Eaters were, on average, unchanged (Figure 2D, Table 6A). Approximately 10% of proteins were differentially detected between the two cell types. However, there were more proteins of macrophage origin enriched on the surface of Non-Eaters compared to Eaters (102 proteins vs. 20 proteins, Figure 2D and Table 6A). This finding suggested that the diversity of native macrophage proteins on the cell surface was decreasing following phagocytosis.

Conversely, over 400 isotopically heavy proteins were enriched on the surface of Eaters compared to Non-Eaters, suggesting that intact proteins synthesized in cancer cells were now on the surface of macrophages that had phagocytosed (Figure 2E, Table 6B). Of the 414 heavy proteins enriched on the surface of Eaters, 395 of these proteins were also detectable on the surface of SW620 cells (Tables 6B and 6C). Proteins abundant on the surface of SW620 cells were more likely to be detected on the surface of Eaters following co-culture (Median = 31%, Extended Data Figure 8). Acquisition of heavy proteins on the cell surface was not observed for macrophages co-cultured with SW620 cancer cells when Eater and Non-Eater populations were not further purified, emphasizing the need to purify the less-abundant Eater population (Extended Data 9, Tables 7A and 7B).

We first confirmed that EpCAM is detectable on the surface of Eater macrophages but not Non-Eaters or naive macrophages in our MS data set (Figure 3A, Table 6B). To validate our findings using orthogonal methods, intact protein integrity was determined by antibody-based flow cytometry (Figure 3B, Extended Data Figure 10). Pharmacologic inhibition of phagocytosis by Cytochalasin D prevented EpCAM detection on the surface of macrophages (+CytoD condition, Figure 3B)^33^. Treatment of macrophages with cancer cell lysate and cancer cell conditioned media (+Lysate) also did not result in detection of EpCAM on the macrophage surface, further suggesting a mechanism dependent on whole cell uptake (Figure 3B). Imaging flow cytometry showed transfer of EpCAM to the cell surface is only observed when cancer cells have entered the macrophage lysosome, as determined by pre-labeling cancer cells with a pH-sensitive rhodamine dye (Figure 3C)^34^. To test if this phagocytosis-dependent surface protein transfer could also occur with other proteins and other cancer cell types, such as blood malignancies, we performed a similar experiment with a B-cell cancer cell line. We observed transfer of CD20 from cancer cell to macrophage that was dependent on whole-cell phagocytosis (Extended Data Figure 11).

**Figure 3.**
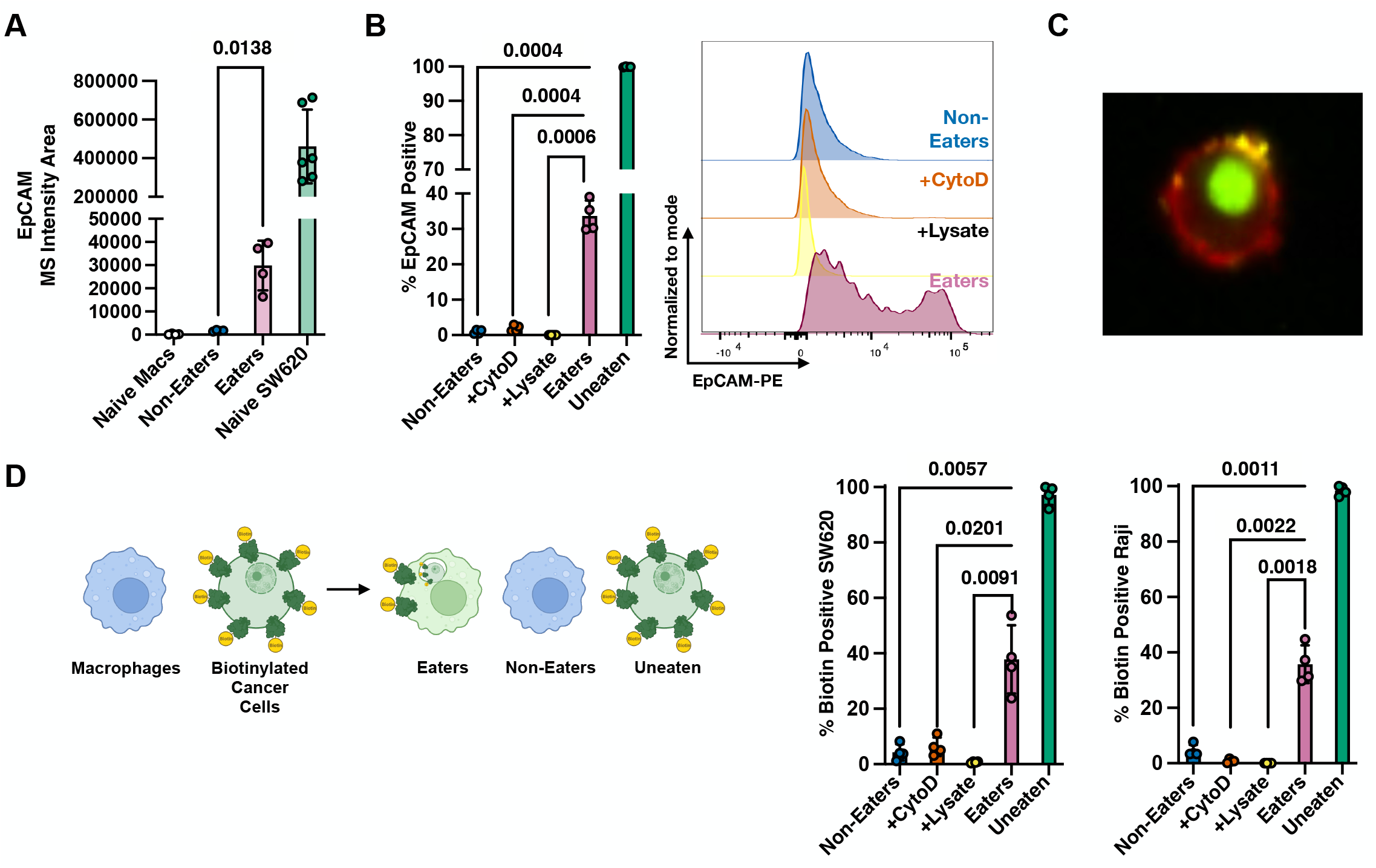
Macrophages that phagocytose adherent and suspension cancer cells incorporate cancer cell-surface proteins onto their own cell surface. A. MS intensity area of heavy-tagged EpCAM detectable on the surface of naive macrophages (Naive Macs), Non-Eaters, Eaters, Naive SW620, or Uneaten cells. n = 2 biological, 2 technical replicates each. B. Flow cytometry analysis of anti-EpCAM staining for cell types of interest from co-culture (Non-Eaters, Eaters and Uneaten SW620. Donor-matched macrophages treated with Cytochalasin D (10 µM) or SW620 conditioned media and lysate were also analyzed. For +Lysate condition, cells were lysed via gentle sonication in their own conditioned culture media prior to addition of macrophages. Values were normalized relative to Uneaten SW620s. n = 4 biological replicates, 2 technical replicates each. C. Representative image showing pHrodo-labeled SW620 cell (green) engulfed by CD11b^+^ macrophage (red), with EpCAM staining (yellow). D. Biotinylated surface protein transfer measurements from macrophages co-cultured with SW620 or Raji cells (2 h). Flow cytometry analysis of anti-Biotin staining for cell types of interest from co-culture (Non-Eaters, Eaters and Uneaten). Donor-matched macrophages treated with Cytochalasin D (10 µM), or cancer cell conditioned media and lysate (+Lysate) were also analyzed. For +Lysate condition, cells were lysed via gentle sonication in their own conditioned culture media prior to addition of macrophages. Values normalized to max on Uneaten cancer cells. n = 2 biological, 2 technical replicates each. Error bars in all figures represent Standard Deviation.

We next developed an assay to test surface protein transfer independent of MS analysis and antigen-specific antibody detection that would also evaluate if the cancer cell surface proteins detected on Eaters had once been on the surface of the cancer cell. Cancer cells were treated with a positively-charged biotin reagent that cannot enter cells and reacts with primary amines on surface proteins. We then co-cultured these biotinylated cancer cells with primary human macrophages (2 h) and tested for the transfer of biotin from cancer cell to macrophage. Biotin transfer was observed for Eaters of both SW620 and Raji cells during phagocytosis (Figure 3D, Extended Data Figure 12). These data demonstrate a mechanism of redeployment of cell surface proteins from the cancer cell surface to the macrophage cell surface following phagocytosis, rather than acquisition of proteins passing through the secretory pathway.

Having confirmed phagocytosis-dependent protein redeployment, we next asked if the transferred proteins are functional. PANTHER analysis of heavy proteins detectable on the surface of Eater macrophages revealed statistical over-representation of transporter proteins compared to a combined macrophage and cancer cell surface reference proteome (Figure 4A)^35,36^. If such proteins were functional following transfer, this could significantly alter the metabolic update of nutrients and metabolites by TAMs, which are known to be metabolically dysregulated^17,37^. By MS and flow cytometry, the active transporter SLC1A5, an established temperature- and pH-sensitive glutamine transporter, was detected as one of the proteins transferred to the Eater surface (Figures 4B and 4C, Table 6B, and Extended Data Figure 13). We generated SLC1A5 KO SW620 and confirmed that SLC1A5 expression in SW620 cells is required for the protein to be detectable on the surface of Eater macrophages following co-culture (Figure 4C and Extended Data Figure 13B).

**Figure 4.**
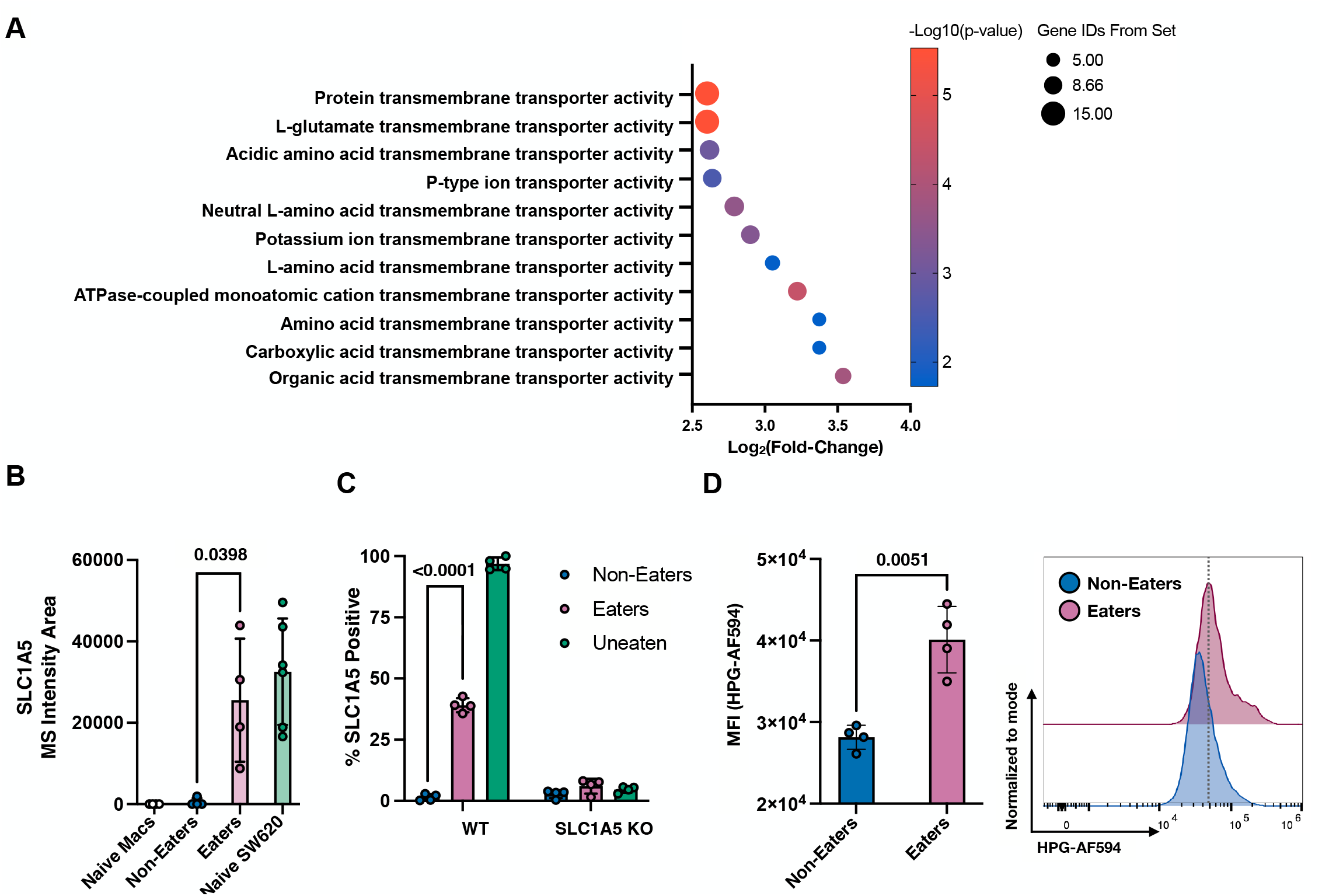
Proteins transferred from cancer cell surface to macrophage cell surface are functional and alter metabolic uptake potential. A. Statistical over-representation analysis of molecular functions of heavy proteins found at least 4-fold increased on the surface of Eater macrophages. Statistical analysis via Fisher’s exact text and correction via Bonferroni. B. MS intensity of SLC1A5 on the surface of Eaters, Non-Eaters, and Uneaten SW620s following co-culture. n = 2 biological, 2 technical replicates each. C. Intracellular flow cytometry analysis of SLC1A5 staining of Eaters, Non-Eaters, and Uneaten SW620 cells following co-culture. Values normalized to max on Uneaten SW620s. Macrophages co-cultured with SLC1A5 KO SW620 cells did not stain for SLC1A5. n = 2 biological, 2 technical replicates each D. Homopropargylglycine (HPG) uptake analysis of Eaters and Non-Eaters following co-culture (2 h). Co-culture samples were incubated with HPG (200 μM, 5 min, 37°C, 5% CO_2_). Transport was quenched upon fixation (2% PFA in PBS, 15 min, 20°C). Cells were permeabilized (0.01% digitonin in PBS, 10 min, 20°C). Internalized HPG was visualized via strain-promoted click chemistry with Az-AF647. n = 2 biological, 2 technical replicates each. All error bars represent standard deviation.

Cancer cells require high levels of glutamine, and cancer cell expression of SLC1A5 is associated with poor prognosis^38,39^. TAMs have previously been shown to exhibit increased glutamine uptake and glutamine metabolism compared to functional macrophages^7,8^. Transfer of additional glutamine transporters to the surface of TAMs, if functional, could explain the etiology of this report. SLC1A5 is one of the few reported transporters that enables uptake of the unnatural amino acid homopropargylglycine (HPG)^40,41^. We went on to demonstrate that Eater macrophages with SLC1A5 on their surface exhibited increased uptake of HPG, a finding consistent with the possibility that proteins redeployed from cancer cell surface to macrophage surface can be functional and are fully incorporated into the macrophage membrane (Figure 4D, Extended Data Figure 14). Importantly, this uptake was sensitive to temperature and glutamine concentration, a characteristic of SLC1A5 function (Extended Data Figure 15).

In this report, we demonstrate that macrophages redeploy proteins from the surface of cancer cells onto their own cell surface during the process of phagocytosis. These proteins are intact and properly oriented, enabling antibody or tag-based recognition. These proteins must also have been surface resident on the cancer cell, as macrophages that have phagocytosed cancer cells pre-labelled with a surface biotin reagent have biotin detectable on their surface. Transporters are statistically over-represented in the dataset of transferred proteins detectable on the surface of macrophages that have phagocytosed, and these proteins appear to be functional.

It is possible that the widespread redeployment of cancer cell surface proteins is related to the known metabolic shifts that have been described for macrophages in the tumor microenvironment. Our functional SLC1A5 data supports this theory. In further support, among the most abundant proteins transferred is the glucose transporter GLUT1 (Extended Data 16, Table 6B)^42^. Other transporters, including additional neutral amino acid transporters as well as the lactate transporter MOT1, are also observed to be redeployed by macrophages following phagocytosis of cancer cells (Extended Data 16, Table 6B). Data generated from our co-culture system is corroborated by human data and two *in vivo* models, the 4T1 breast cancer model and the myeloproliferative neoplasm JAK2 V617F model. In both models the presence of non-native macrophage proteins detectable on the surface of macrophages is associated with intracellular protein content diagnostic of phagocytosis. Proteins detectable from whole cell MS studies are associated with cell compartments beyond the cell membrane, and mRNA from these cells is detectable in transcriptomic studies, further supporting a mechanism of whole cell uptake and not trogocytosis is responsible for the protein redeployment observed in our systems. Moreover, redeployment is not observed upon inhibition of phagocytosis or incubation of macrophages with cancer cell lysates and conditioned media. MS analysis of our isotopically labelled co-culture system and surface biotinylation studies suggest a mechanism of surface protein redeployment, rather than the acquisition of cancer cell mRNA. Further work will be required to understand the precise mechanism of protein redeployment so that it can be perturbed both to understand the mechanistic intricacies of the process and to test its impact on innate immune control of cancer.

The work described suggests a novel model in TAM etiology. It is not that TAMs are intrinsically poor phagocytes, rather it is that they have phagocytosed so many cancer cells that they have reengineered their cell surface, redeploying proteins that increase metabolic uptake while diluting out or losing surface proteins that promote phagocytosis. It has been well established that ligand-receptor interactions modulating phagocytosis are often avidity-driven events, where protein concentration is critical. Notably, this redeployment mechanism is transcriptionally silent, significantly altering the function of a TAM while not requiring gene transcription or protein synthesis. This hypothesis is consistent with our previous finding that macrophages have low rates of translation^30^. It is possible that this protein redeployment mechanism extends to the programmed cell removal of apoptotic and necrotic cells, and this question is currently under investigation in our laboratory. While our macrophage co-culture system focuses on live-cell phagocytosis, we cannot exclude the possibility that some protein redeployment events observed in our *in vivo* models are occurring through necrotic or apoptotic cell uptake mechanisms. Nevertheless, our work undoubtedly demonstrates that redeployment of surface proteins of cancer cell origin onto the surface of macrophages has the potential to alter macrophage function and metabolic state.

## Supporting information

Volk2024_ExtendedData

Volk2024_Methods

SourceData

Table4_RNA_EatersNon

Table5_WholeCellMS_EatersNon

Table6_SurfaceMS_EatersNon

Table7_SurfaceMS_CoCulture

Table8_Antibodies

Tables2+3_MS_Spleen

## Acknowledgements

We thank members of the Zaro, Goodarzi, Fidler, Maker and Ramani laboratories for advice and discussions. We thank the Gladstone Flow Cytometry Core and J. Srivastava for her invaluable work in maintaining the facility. We thank H. Hang, M. Pratt, H. Madhani, J. Moslehi, C. Devereux, and J. Tenthorey for careful reading of the manuscript. The research reported in this publication was supported by Arnold and Mabel Beckman Foundation (to B.W.Z), the Program in Breakthrough Biomedical Research, partially funded by the Sandler Foundation (to B.W.Z), the UCSF Discovery Fellowship (to R.F.V, to S.W.C.), the Achievement Rewards for College Scientists Scholarship (to R.F.V), the UCSF Pharmacy Graduate Program Fellowship (to R.F.V), the UCSF J. Michael Bishop Fellowship (R.F.V), the National Science Foundation Graduate Research Fellowship under Grant No. 1650113 (to S.W.C.). The funders had no role in study design, data collection and analysis, decision to publish, or preparation of the manuscript.

## Contributions

R.F.V. conceived and designed experiments, performed experiments, analyzed data and prepared the manuscript. S.W.C. conceived and designed experiments, performed experiments, analyzed data and edited the manuscript. A.C.C. performed experiments and edited the manuscript. B.Z., N.M., and I.I. performed experiments, analyzed data, and edited the manuscript. H.L. prepared mouse samples and edited the manuscript. S.T. and T.P. processed primary human tissues and edited the manuscript. V.R. conceived and designed experiments and supervised the research. A.V.M. supervised the collection and processing of primary human tissues and edited the manuscript. T.F. conceived and designed experiments, performed experiments, and edited the manuscript. H.G. conceived and designed experiments, analyzed data, and supervised the research. B.W.Z. conceived and designed experiments, analyzed data, supervised the research, and wrote the manuscript.

