## Supplementary material for "Macrophages redeploy functional cancer cell surface proteins following phagocytosis": Volk2024_ExtendedData

**Volk et al Extended Data Files**

Table List
Extended Data Figures 1-16

**Volk et al Table List**

Table 1. 4T1 Mass Spectrometry Data

Table 2. Mouse Spleen Mass Spectrometry Data

Table 3. Mouse Red Blood Cell Mass Spectrometry Data

Table 4. RNA-Sequencing of Eaters and Non-Eaters

Table 5A. Whole Cell Mass Spectrometry Data of Eaters and Non-Eaters – Light

Table 5B. Whole Cell Mass Spectrometry Data of Eaters and Non-Eaters – Heavy

Table 5C. Whole Cell Mass Spectrometry Data of Eaters and Non-Eaters – Heavy + Light

Table 6A. Surface Enrichment Mass Spectrometry Data of Eaters and Non-Eaters – Light

Table 6B. Surface Enrichment Mass Spectrometry Data of Eaters and Non-Eaters – Heavy

Table 6C. Surface Enrichment Mass Spectrometry Data of SW620 Cells

Table 7A. Surface Enrichment Mass Spectrometry Data of Co-Cultured Macrophages – Light

Table 7B. Surface Enrichment Mass Spectrometry Data of Co-Cultured Macrophages – Heavy

Table 8. Antibodies Used

Table 9. Source Data

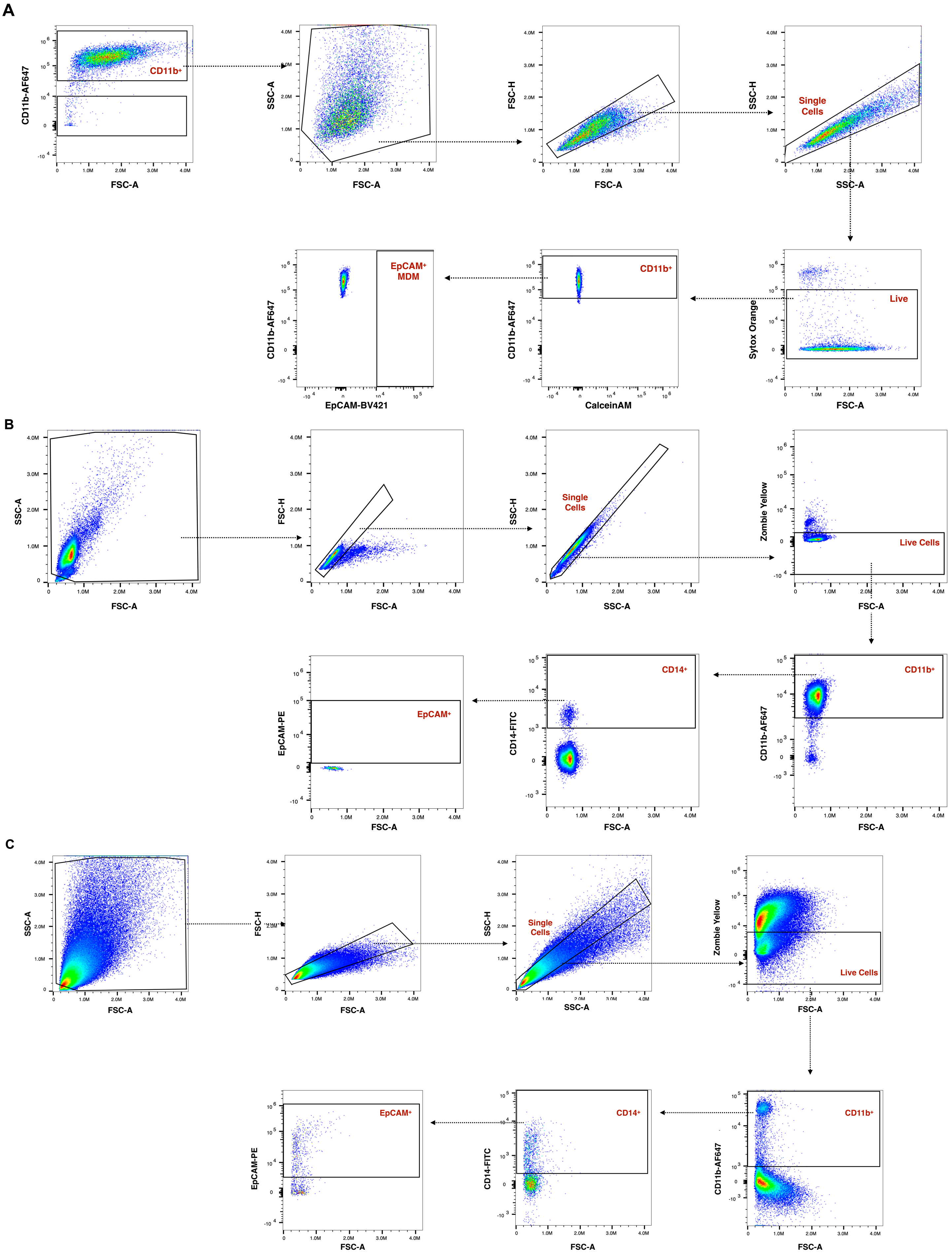

A
CD11b-AF647
FSC-A
CD11b+
SSC-A
FSC-H
SSC-H
Single Cells
Sytox Orange
Live
CalceinAM
EpCAM-BV421
EpCAM+ MDM
B
Zombie Yellow
Live Cells
CD14-FITC
CD14+
EpCAM-PE
EpCAM+
C

**Extended Data Figure 1. EpCAM is detectable on the surface of macrophages from primary human tumors and not in the periphery.** Gating strategy to evaluate EpCAM positivity on tumor associated macrophages (TAMs, C) compared to monocyte-derived macrophages (A) and paired peripheral blood macrophages (B). After excluding debris, doublets and dead cells, live macrophages were identified as $CD11b^{+}$ $CD14^{+}$ in paired TAMs and peripheral blood samples. Live macrophages were identified as $CD11b^{+}$ from monocyte-derived macrophages. Cells were further separated dependent on the presence of EpCAM on their surface. Plots are representative of 3 biological replicates.

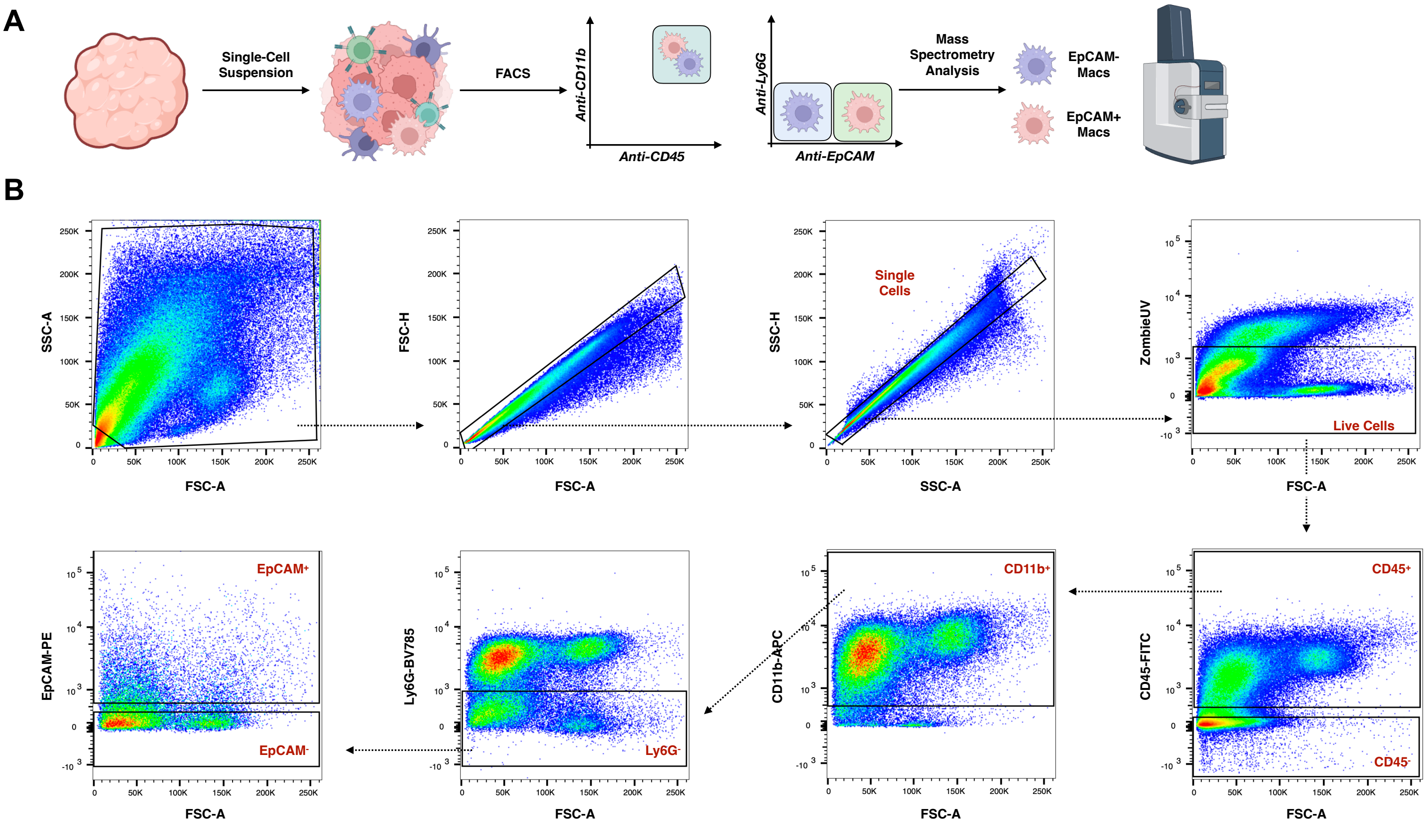


**Extended Data Figure 2. EpCAM is detectable on the surface of tumor associated macrophages in a 4T1 model of breast cancer.** A. Experimental workflow. 4T1 tumors were excised and processed to generate single-cell suspensions. Cell suspensions were stained with desired flow cytometry-compatible antibodies and subjected to FACS in order to purify $EpCAM^{+}$ and $EpCAM^{-}$ tumor associated macrophages. Purified populations were processed for mass spectrometry analysis to compare proteins detectable in each cell population. B. Gating strategy for tumor associated macrophages. After excluding debris, doublets and dead cells, live macrophages were identified as $CD45^{+}CD11b^{+}Ly6G^{-}$. Cells were then further separated dependent on the presence of EpCAM on their surface. Plots are representative of 6 biological replicates.

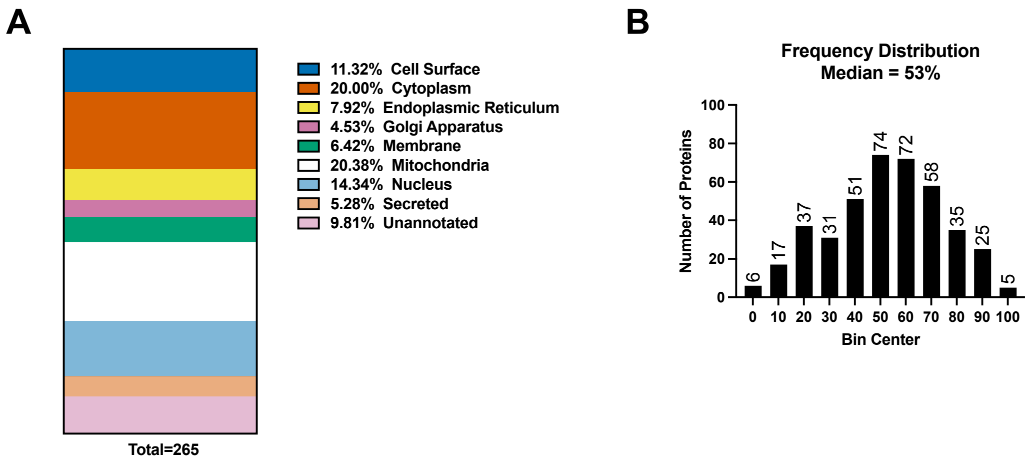


**Extended Data Figure 3. Characterization of 4T1 proteins detectable in EpCAM⁺ macrophages.** A. Cellular localization by Uniprot annotation of proteins likely originating from EpCAM-expressing 4T1 cells detectable in EpCAM⁺ macrophages. B. Abundance distribution of 4T1 cell proteins detectable in EpCAM⁺ macrophages with respect to their abundance in 4T1 cell proteomes reveals no bias in detection towards abundant proteins. Bin centers are annotated by percentile abundance. For example, Bin 0 represents the top 10% most abundant proteins in 4T1 cells as detected by MS.

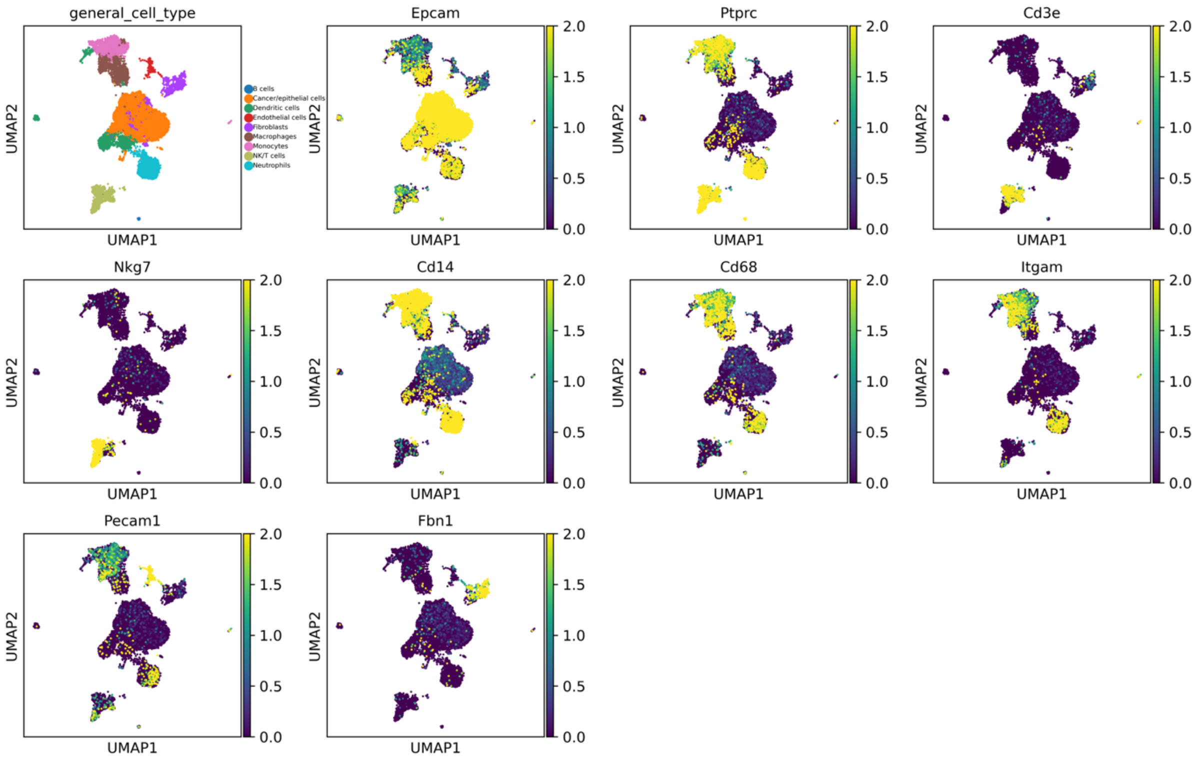


**Extended Data Figure 4. EPCAM is detectable within macrophages in single cell sequencing of 4T1 tumors.** Cluster annotation using automatic cell type annotation and expression of cell type markers across isolated cells, including monocyte/macrophage specific markers and EpCAM. Data is derived from GEO Accession: GSE230641.

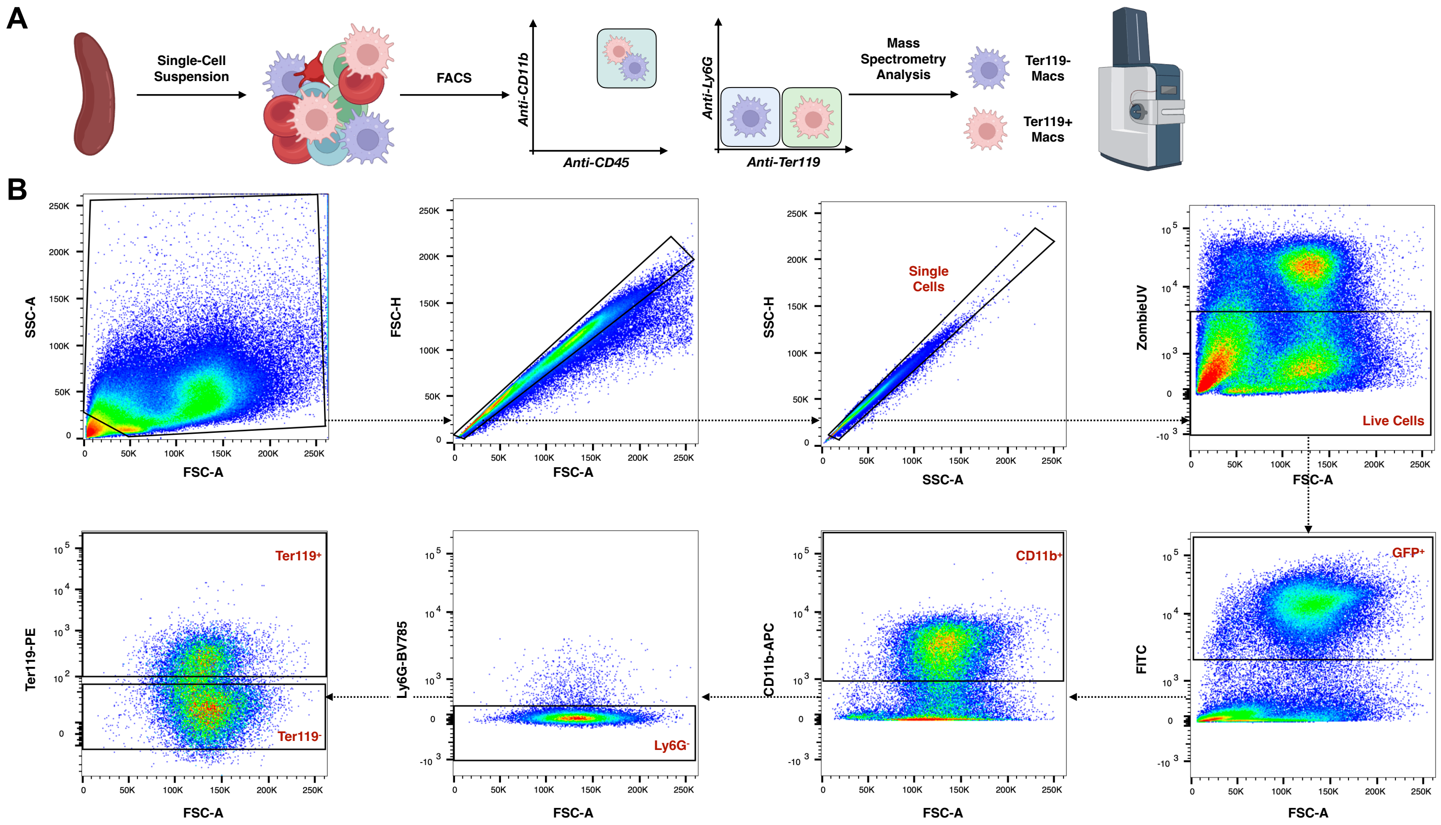


**Extended Data Figure 5. Ter119 is detectable on the surface of macrophages in the spleen.** A. Experimental workflow. Spleens were excised and processed to generate single-cell suspensions. Cell suspensions were stained with desired flow cytometry-compatible antibodies and subjected to FACS to purify Ter119$^{+}$ and Ter119$^{-}$ macrophages. Purified populations were processed for mass spectrometry analysis to compare proteins detectable in each cell population. B. Gating strategy for macrophages in the spleen. After excluding debris, doublets and dead cells, live macrophages were identified as GFP$^{+}$CD11b$^{+}$Ly6G$^{-}$. Cells were then further separated dependent on the presence of Ter119 on their surface. Plots are representative of 8 biological replicates.

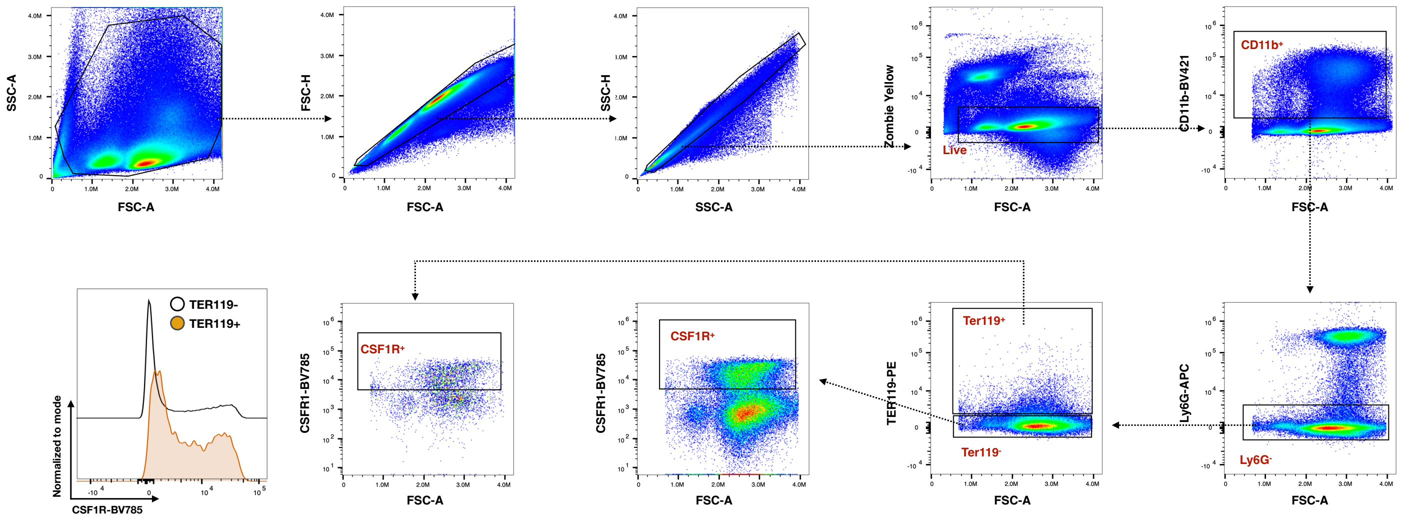


**Extended Data Figure 6. CSF1R levels are different on the surface of Ter119$^{+}$ and Ter119$^{-}$ macrophages in the spleen.** Gating strategy for macrophages in the spleen. After excluding debris, doublets and dead cells, live macrophages were identified as GFP$^{+}$CD11b$^{+}$Ly6G$^{-}$. Cells were then further separated dependent on the presence of Ter119 on their surface and then CSF1R. Plots are representative of 4 biological replicates.

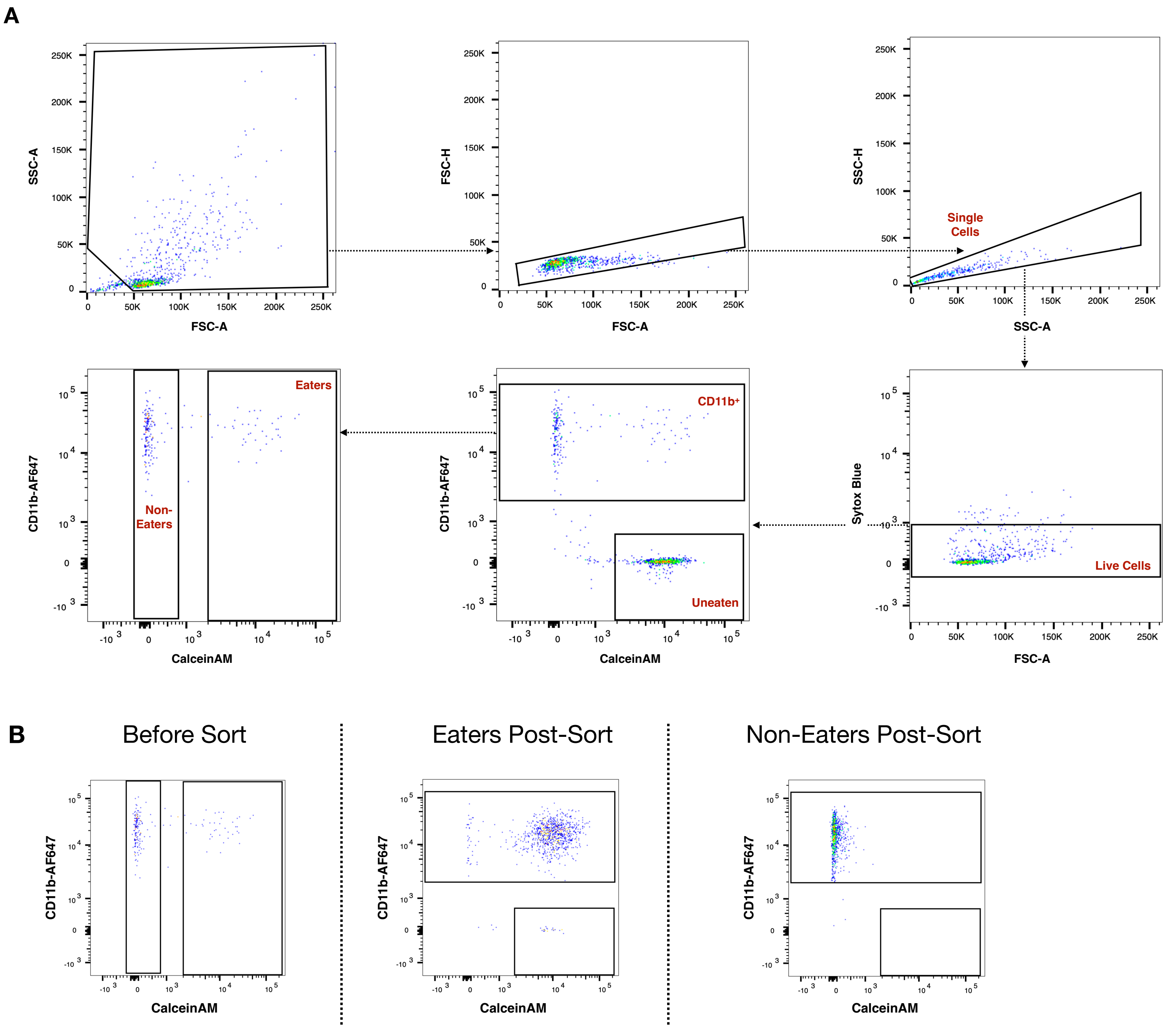


**Extended Data Figure 7. A macrophage phagocytosis co-culture assay.** A. Gating strategy for a cell culture-based phagocytosis assay with mass spectrometry readout. Following removal of debris, doublets, and dead cells, Non-Eaters were defined as $CD11b^{+}FITC^{-}$ and Eaters as $CD11b^{+}FITC^{+}$. B. Representative purity for cell populations subjected to mass spectrometry analysis (Eaters and Non-Eaters). Plots are representative of data from 2 biological replicates.

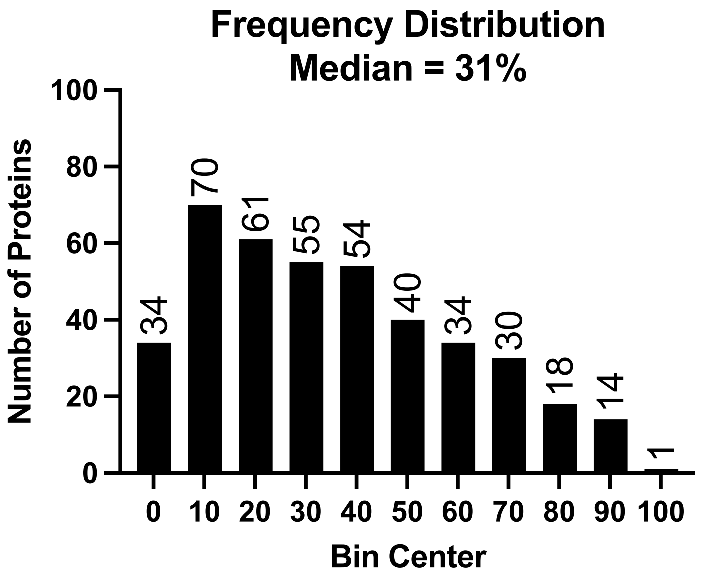


**Extended Data Figure 8. Detection of surface of Eaters correlates with SW620 cancer cell surface abundance.** Frequency distribution of heavy proteins detectable on the surface of Eaters with respect to abundance on the surface of SW620 cells. Bin centers are annotated by percentile abundance. For example, Bin 0 represents the top 10% most abundant proteins in SW620 cells as detected by MS.

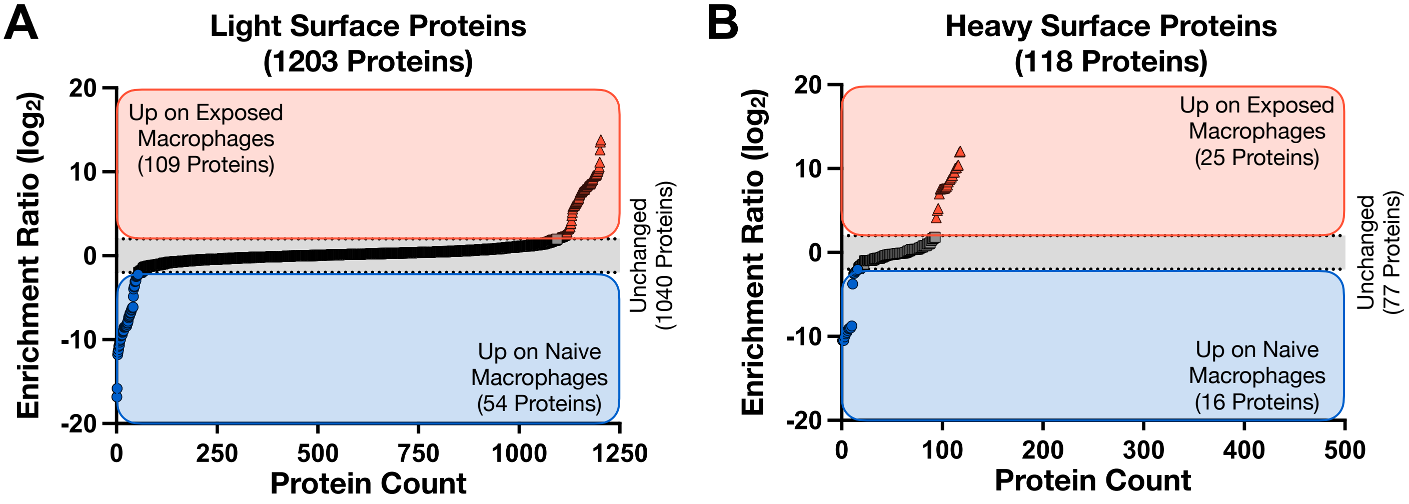


**Extended Data Figure 9. Mass spectrometry analysis of macrophages co-cultured with cancer cells does not reveal the heavy surface protein signature observed when Eaters and Non-Eaters are purified.** A. Relative abundance of light surface proteins detectable on macrophages co-cultured with cancer cells compared to naive macrophages. B. Relative abundance of heavy surface proteins detectable on macrophages co-cultured with cancer cells compared to naive macrophages. X-axes were selected to enable direct comparison to Figures 2D and 2E.

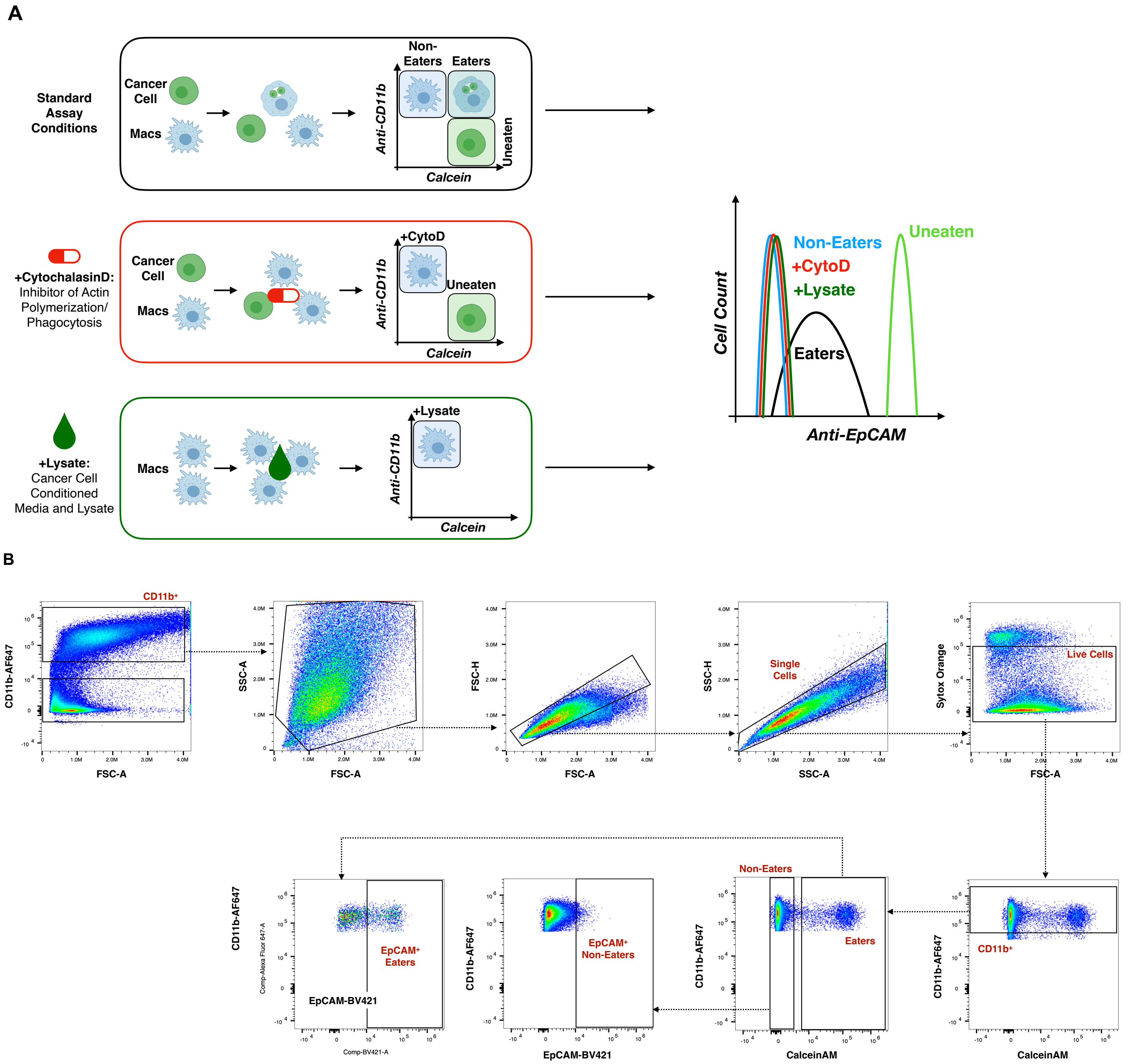


**Extended Data Figure 10. EpCAM is detectable on the surface of Eater macrophages.** A. Experimental workflow to evaluate EpCAM expression on the surface of macrophages is dependent on phagocytosis. Macrophages were co-cultured with cancer cells pre-stained with the intracellular viability dye CalceinAM. To evaluate dependence of phagocytosis, the co-culture system was treated with Cytochalasin D, which inhibits actin polymerization, resulting in no 'Eater' macrophages. To evaluate dependence on intact cell interactions, a cell lysate and conditioned media mixture was generated from cancer cells gently lysed by sonication in their conditioned media. In both +CytoD and +Lysate conditions, all macrophages were evaluated for EpCAM positivity. B. Gating strategy to evaluate EpCAM positivity on Eater and Non-Eater macrophages following co-culture with SW620 cancer cells. After excluding debris, doublets and dead cells, Non-Eaters as $CD11b^{+}FITC^{-}$ and Eaters as $CD11b^{+}FITC^{+}$. Cells were then further separated dependent on the presence of EpCAM on their surface. Plots are representative of 4 biological, 8 technical replicates.

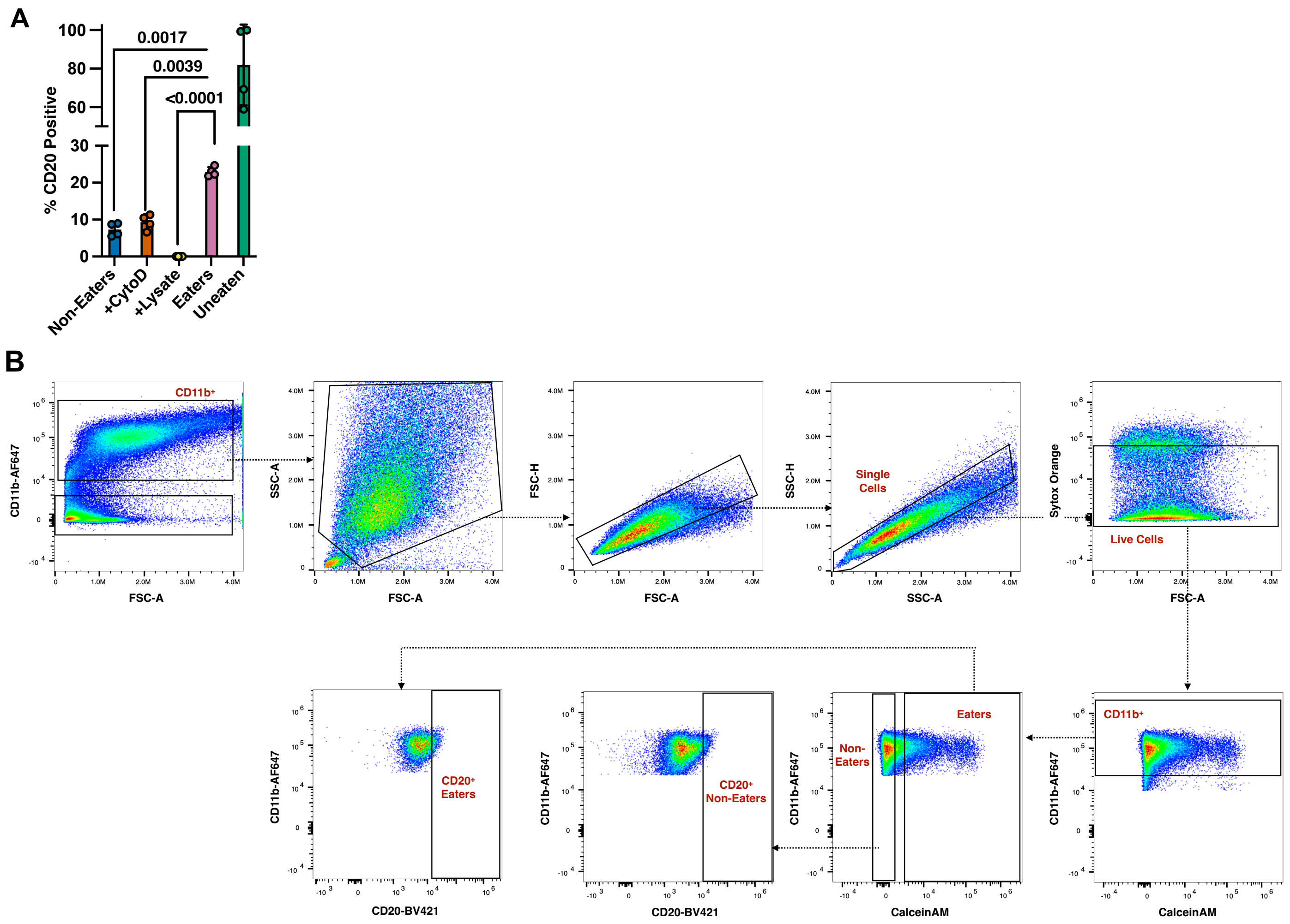


**Extended Data Figure 11. CD20 is detectable on the surface of Eater macrophages.** A. Flow cytometry analysis of anti-CD20 staining for cell types of interest from co-culture (Non-Eaters, Eaters and Uneaten Raji. Donor-matched macrophages treated with Cytochalasin D (10 µM) or Raji conditioned media and lysate were also analyzed. For +Lysate condition, cells were lysed via gentle sonication in their own conditioned culture media prior to addition of macrophages. Values normalized to max on Uneaten Rajis. N = 2 donors, 2 experimental replicates per donor. Error bars denote S.D. B. Gating strategy to evaluate EpCAM positivity on Eater and Non-Eater macrophages following co-culture with Raji cancer cells. After excluding debris, doublets and dead cells, Non-Eaters were identified as $CD11b^{+}FITC^{-}$ and Eaters as $CD11b^{+}FITC^{+}$. Cells were then further separated dependent on the presence of CD20 on their surface. Plots are representative of 2 biological, 4 technical replicates.

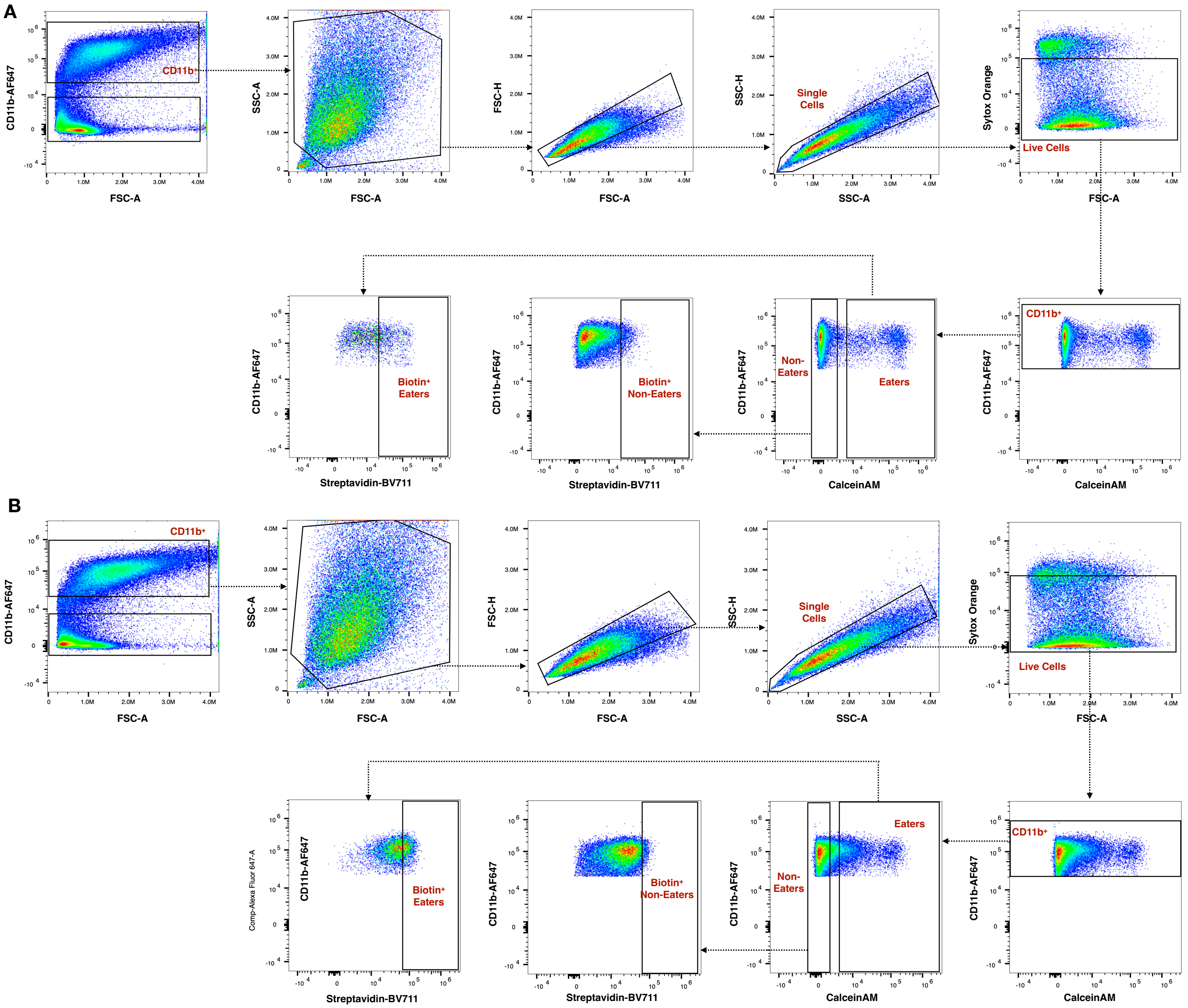


**Extended Data Figure 12. Biotin is detectable on the surface of Eater macrophages.** Gating strategy to evaluate Biotin positivity on Eater and Non-Eater macrophages following co-culture with calcein-stained SW620 (A) or Raji cancer cells (B). After excluding debris, doublets and dead cells, Non-Eaters as $CD11b^{+}FITC^{-}$ and Eaters as $CD11b^{+}FITC^{+}$. Cells were then further separated dependent on the presence of biotin on their surface. Plots are representative of 3 biological, 6 technical replicates.

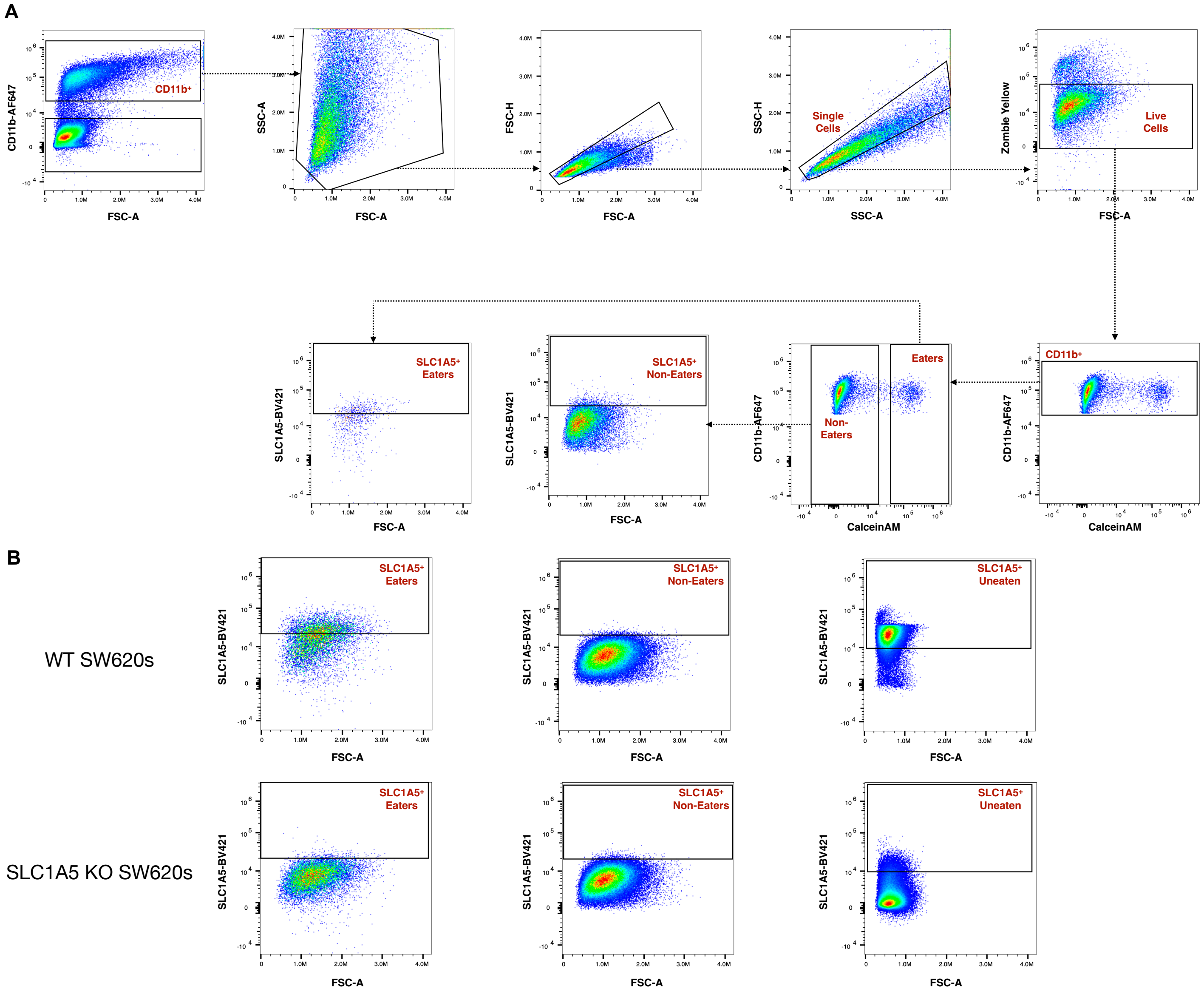


**Extended Data Figure 13. SLC1A5 is detectable on the surface of Eater macrophages following phagocytosis.** A. Gating strategy to evaluate SLC1A5 positivity on Eater and Non-Eater macrophages following co-culture with wild-type (WT) or SLC1A5 KO SW620 cells. After excluding debris, doublets and dead cells, Non-Eaters were defined as CD11b$^{+}$FITC$^{-}$ and Eaters as CD11b$^{+}$FITC$^{+}$. Cells were then further separated dependent on the presence of SLC1A5 on their surface. Plots are representative of 4 biological, 8 technical replicates. B. Comparison of SLC1A5 staining of Eaters and Non-Eaters following co-culture with WT SW620 cells or SLC1A5 KO SW620 cells.

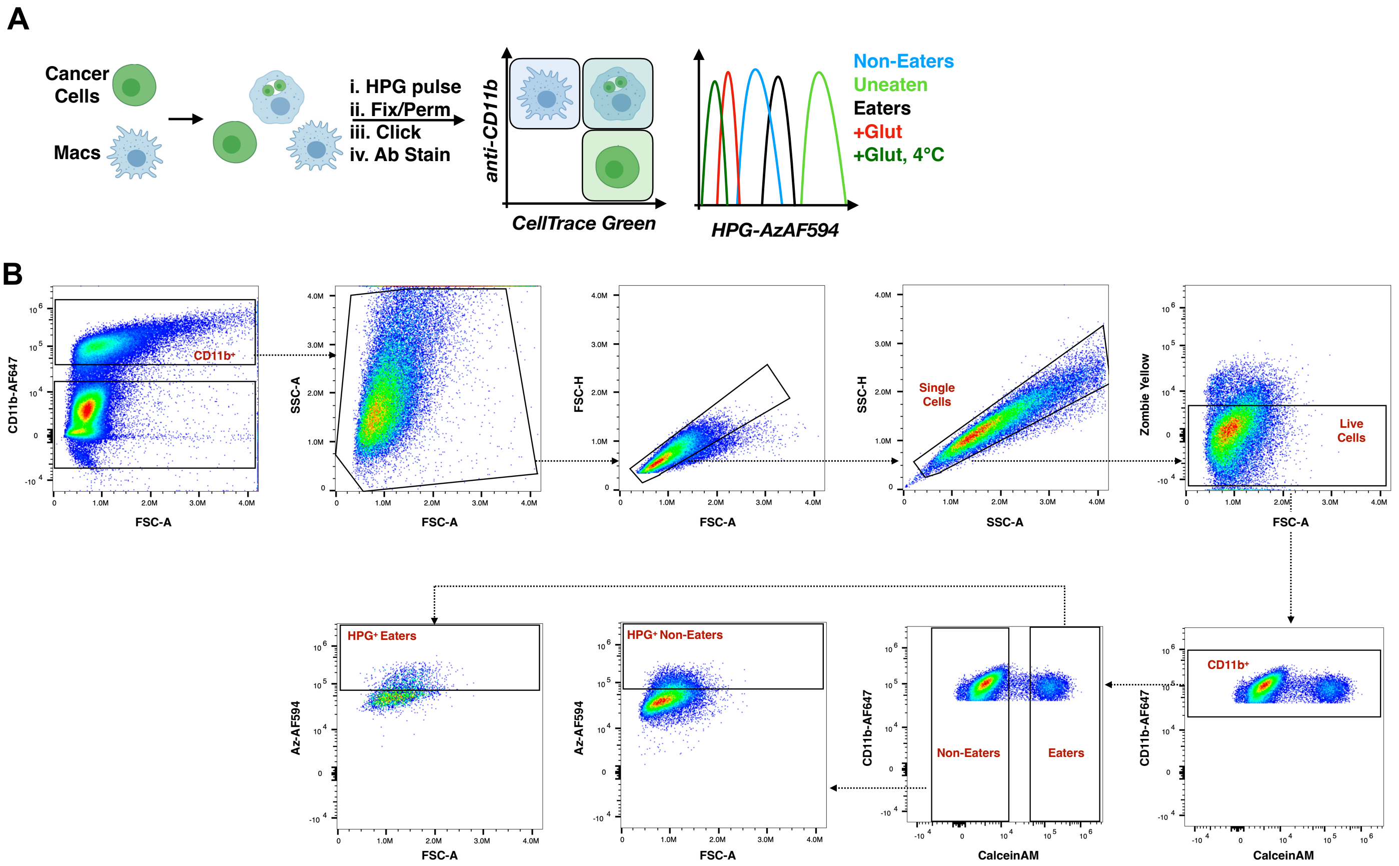


**Extended Data Figure 14. HPG is taken up by Eater macrophages more than Non-Eater macrophages.** A. Experimental workflow. SW520 cancer cells pre-stained with an intracellular viability dye and macrophages were co-cultured for 2h after which time the cell mixture was pulsed with homoproprargylglycine (HPG) for 5 min. Cells were washed to remove excess HPG prior to fixation and permeabilization. The HPG-labelled co-culture mixture was subjected to click chemistry with Azide-AF594 and stained for CD11b and viability prior to flow cytometry analysis. B. Gating strategy to evaluate HPG uptake by Eater and Non-Eater macrophages following co-culture with SW620 cells. After excluding debris, doublets and dead cells, Non-Eaters were defined as CD11b$^{+}$FITC$^{-}$ and Eaters as CD11b$^{+}$FITC$^{+}$. Cells were then further separated dependent on the presence of internalized HPG measured by clicked on Az-AF594.

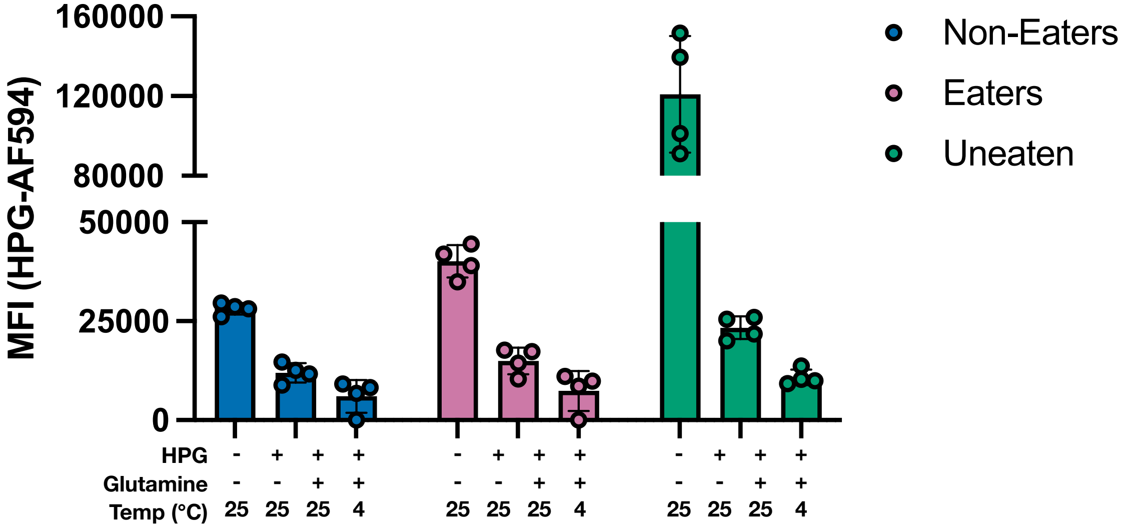


**Extended Data Figure 15. The unnatural amino acid HPG is taken up by Eaters more efficiently Non-Eaters.** Following co-incubation (2 h), samples were incubated with 200 µM HPG for 5 mins at 37℃, 5% $CO_2$. Transport was quenched by fixation (2% PFA in PBS, 15 min). Cells were then permeabilized (0.01% digitonin in PBS, 10 min). Internalized HPG was visualized following click chemistry with Az-AF647. Transport was competed with the addition of 2mM glutamine at 37℃ or 4℃. N = 2 donors, two technical replicates per donor. Error bars represent standard deviation.

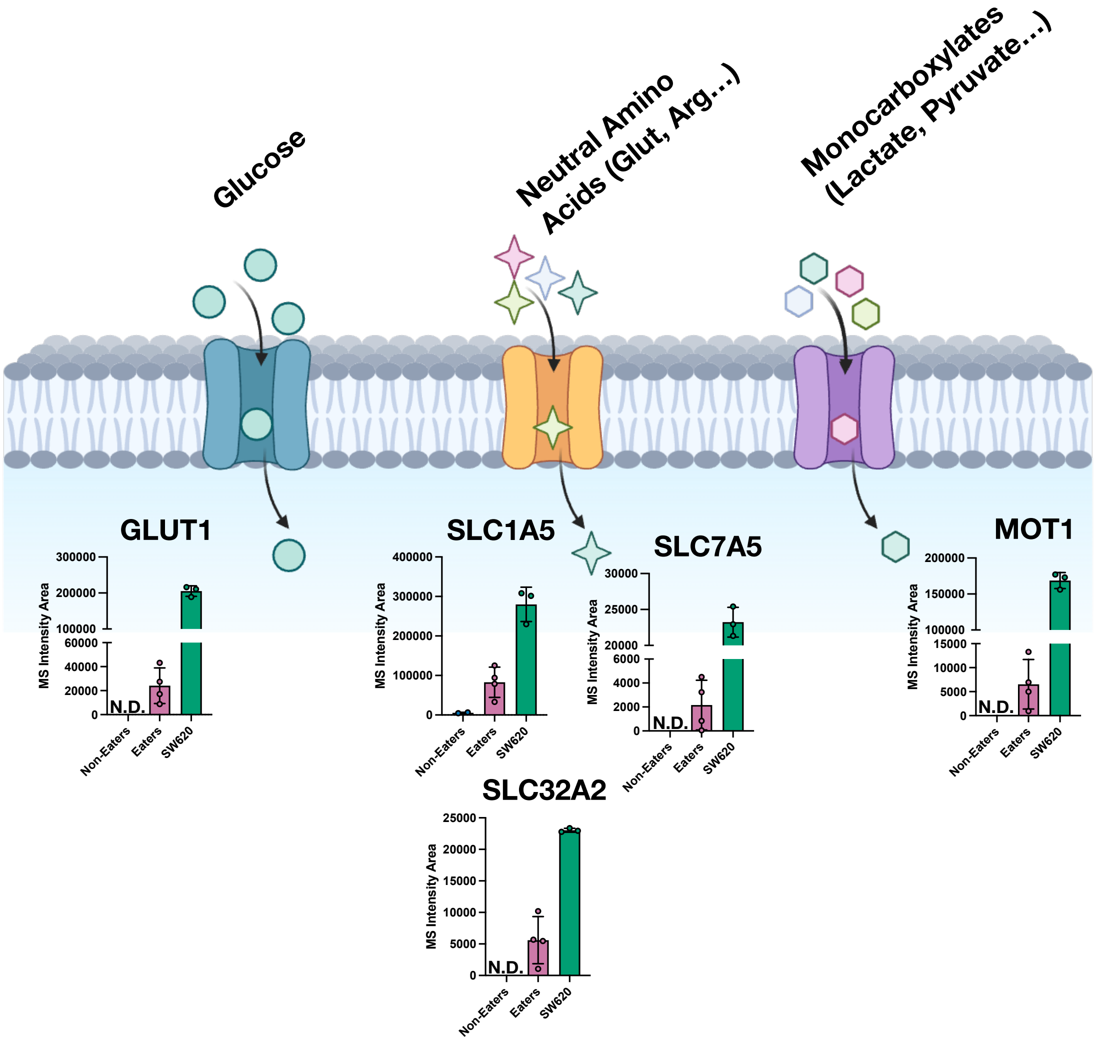


**Extended Data Figure 16. Nutrient transporters are transferred to the surface of Eaters during phagocytosis.** MS intensity of 5 nutrient transporters on the surface of Eaters and Non-Eaters following co-culture with SW620 colorectal cancer cells. N = 2 donors, 2 technical replicates per donor. N.D. denotes not detected in any of the 4 replicates. Error bars represent standard deviation.
