## Supplementary material for "Macrophages redeploy functional cancer cell surface proteins following phagocytosis": Volk2024_Methods

**General Cell Culture**

All cell lines were purchased from American Type Culture Collection (ATCC). SW620 cells were cultured in Hyclone DMEM (Cytiva, SH3008101) supplemented with 2mM GlutaMAX (Gibco, 35-050-061), 10% Fetal Bovine Serum (FBS, Cytiva, SH3008803), 100U/mL penicillin, and 100µg/mL streptomycin (VWR, 97063-708). SW620 were maintained in TC-Treated Culture Dishes (Corning, 877222). Raji cells were cultured in RPM-1640 (ATCC, 30-2001) supplemental with 10% Fetal Bovine Serum (FBS, Cytiva, SH3008803), 100U/mL penicillin, and 100µg/mL streptomycin (VWR, 97063-708) in a Suspension Culture Flask (Greiner, 690195). All cells were grown in a 37°C, 5% CO2 incubator.

**Culture of SILAC Heavy Cells**

SW620 were cultured in DMEM for SILAC (Thermo Scientific, PI88364) supplemented with 100µg/mL L-Arginine·HCl, 100µg/mL L-Lysine·2HCl (Cambridge Isotope Laboratories Inc., CNLM-539-H-1, CNLM-291-H-1), 10% Dialyzed Fetal Bovine Serum (Thermo Scientific, 26400044), 100U/mL penicillin, and 100µg/mL streptomycin (VWR). Cells were grown in a 37°C, 5% CO2 incubator. SW620 were passaged in media containing heavy amino acids 6 times to allow for complete incorporation. Level of incorporation was evaluated by LC-MS (Bruker) prior to implementation in experiments.

**CRISPR-Cas9 Knockout of SLC1A5 in SW620 Cells**

$SLC1A5^{-/-}$ cells were generated by CRISPR-Cas9 mediated genome editing. Guide RNA (5'-ACTCTACCACCTATGAAGAG-3') was complexed with tracrRNA (IDT, 1072532) before being added to Cas9 nuclease (IDT, 10008100). The resulting RNP and electroporation enhancer (IDT, 1075915) were nucleofected into SW620 cells using the Lonza 4D-Nucleofector and SF Cell Line 4D-Nucleofector X Kit (Lonza, V4XC-2032). Genomic editing was confirmed using Synthego ICE analysis and loss of SLC1A5 protein was confirmed by flow. Single cell clones were isolated by limiting dilution.

**Animal Care and Experiments**

All animal studied were performed in accordance with the institutional animal care and use committee of the University of California, San Francisco.

<u>Jak2V617F Mice</u>

All experiments utilized bone marrow transplantation (BMT), with donor mice sex-matched to recipients. Donor mice were 6–14 weeks old and recipient C57Bl6J wild type mice were 8–12 weeks old. All mice used for these studies were on a C57Bl/6J background and were housed at UCSF in a dedicated pathogen-free facility under standard conditions of temperature with a 12-h light/dark cycle and food available ad lib. Jak2V617F conditional knock-ins have been previously described[1] and were induced with an Mx1-Cre (Jackson Lab, B6.Cg-Tg(Mx1-cre)1Cgn/J, 003556). C57Bl6J mice were lethally irradiated with 9.5gy, then transplanted with littermate control or Mx1-Cre+Jak2V617F bone marrow. Mice were allowed to recover for 6 weeks, then spleens were harvested for single cell suspensions as described below[2].

<u>Metastatic 4T1 Mouse Mammary Carcinoma</u>

8–10-week-old age-matched female Balb/cJ (000651) mice were injected bilaterally into mammary fat pads. Each MFP received 500,000 cells resuspended in 50 ul of 1:1 Matrigel (Corning, CB-40230) and PBS. Tumors were monitored for 2-3 weeks, or until they reached a size of ~ 1000 mm3, before dissection. All mice were purchased from Jackson Laboratory.

**scRNASeq Analysis**

Single cell sequencing data of 4T1 mammory tumors was acquired from GEO Accession: GSE230641.

**Flow Cytometry-Based Phagocytosis Assay**

Evaluation of surface protein transfer from calcein stained cancer cells to monocyte derived macrophages was performed by modifying methods outlined in Barkal *et al.* 2019[3]. All actions were performed with cells on ice and centrifuged at 4°C unless stated otherwise. In brief, immortalized cancer cells were stained with Calcein AM (Thermo Fisher Scientific, C34852) following vendor specifications. Cells were washed 2x with 10mLs of DPBS and resuspended at 1 million cells per mL. Calcein AM reagent aliquot was resuspended in DMSO and then diluted to 1mM working stock. 2μL of working stock was added for each mL of cells being stained and incubated at RT for 20mins. Monocyte-dervied macrophages were lifted from plates by washing 3x with DPBS and then incubating with TrypLE for 15mins at 37°C. Following incubation, 5mLs of RPMI Complete which consists of Hyclone RPMI 1640 (Cytiva, SH30255.01) supplemented with 2mM GlutaMAX (Gibco, 35-050-061), 10% Fetal Bovine Serum (FBS, Cytiva, SH3008803), 100U/mL penicillin, and 100μg/mL streptomycin (VWR, 97063-708) was added. A 1mL pipette was used to wash cells from plate before transferring to a falcon tube. All cells were washed 3x with 10mLs DPBS prior to combining in a 100,000:50,000 (cancer cells:macrophage) ratio. Combined cells were centrifuged at 500xg for 5mins and resuspended in IMDM supplemented with GlutaMax and 1ug/mL CD47 blocking reagent (Magrolimab, SelleckChem, A2036) at 750,000 cells (cancer cells+macrophages) per mL. In relevant samples, cytochalasin D (Sigma, C8273) was added following resuspension at a final concentration of 10μM. For macrophages treated with lysate, the 100,000 cancer cells were lysed into their culture medium via probe sonication (1sec on/off, 30 seconds total) and then passed through a 0.45μm syringe filter (VWR, 22023) prior to introduction to 50,000 macrophages. Combined cells were incubated in a 96-Well Clear Ultra Low Attachment U-Bottom Microplates (Corning, 7201680), adding 200μL per well. Assay was incubated for 2hrs in an incubator at 37°C with 5% CO2 before moving to ice to prevent further phagocytosis. Cells were transferred from 96-Well plate to Falcon Round-Bottom Polystyrene Test Tubes with Cell Strainer Snap Cap (Fisher, 877123) passing them through the filter cap to resolve clumping. Samples were centrifuged at 500xg for 5mins, and co-incubation media aspirated before washing 1x with 1mL of FACS Buffer (DPBS, 2% Fetal Bovine Serum). Cells were then resuspended in 100μL FACS buffer and stained with surface antibodies on ice for 30mins at concentrations listed in Supplemental Table 8. After staining, cells were washed with 1mL of FACS buffer and then resuspended in FACS buffer with Sytox Orange (1:10,000, Thermo Fisher Scientific, S34861) viability dye and incubated 10mins at RT prior to analysis on Cytek Aurora Spectral Flow Cytometer.

**Imagestream**

Phagocytosis assay was set up as described above, but SW620s were stained with the pH sensitive dye pHrodo Green STP Ester (Thermo Fisher Scientific, P35369) according to manufacturer's instructions. Following lifting of SW620s from culture plates, cells were washed once with DPBS, and then resuspended in DPBS at 1million/mL. pHrodo Green STP was resuspended in DMSO at 1mg/mL and 1μL of dye was added for each 1mL of cells. Sample was gently agitated to mix and incubated at RT for 30mins. Cells were then washed twice with DPBS supplemented with 1% BSA (VWR, 7061-420). Cells then proceeded through co-incubation with macrophages and antibody labeling as described. Multispectral image analysis was performed on an Imaging Flow Cytometer (Cytek, Amnis). Laser settings were 488nm (10mW), 561nm (100mW), 642nm (150mW), and 785nm (0.5mW). Following collection, images were analyzed using ImageStream Data Analysis and Exploration Software (IDEAS, Version 6.2.187.0, Amnis).

**FACS Sorting for Mass Spectrometry Analysis**

Fluorescence-Activated Cell Sorting assay was set up as described above using calcein stained isotopically tagged SW620 cancer cells. After co-incubation of SW620s and macrophages in 96-well plate, cells were stained with Anti-CD11b according to Supplemental Table 8. Cells were washed with 1mL FACS buffer and then resuspended in FACS buffer with Sytox Blue (1:1000, Thermo Fisher Scientific, S34857). Samples were sorted on a BD Aria III into FACS tubes coated with FACS buffer and cooled to 4°C. After collection, cells were centrifuged at 500xg for 5mins and immediately transferred to Protein LoBind

Microcentrifuge Tubes (Fisher, 13698794) with DPBS to reduce carry-over of contaminating FBS proteins. Cells were then washed 3x with 1mL DPBS and subjected to surface biotinylation or snap freezing and storage at -80C.

**Preparation of single cell suspension from mouse spleen**

Excised spleens were placed in a petri dish with 5mLs of RPMI (Cytiva) and a razor blade was used to cut the organ into smaller pieces. All the following steps were performed at 4°C or on ice unless stated otherwise. The broken-down tissue was transferred to a 40μm Cell Strainer (Falcon, 08-771-1) placed in a 50mL Falcon tube and pushed through using a double edge Cell Lifter (VWR, 76036-006). RPMI was continuously added in 5mL aliquots to support pushing cells through strainer to generate single cell suspension. After tissue was fully through strainer, volume was adjusted to 50mLs with RPMI and then cells were centrifuged at 500xg for 5mins. The samples were resuspended in 15mLs of RBC Lysis Buffer (Thermo Fisher, 00-4333-57) and incubated for 5mins. FACS buffer was added up to 50mLs and centrifugation was repeated. The filtration step through the cell strainer was repeated, substituting RPMI for FACS buffer. After centrifugation, cells were either resuspended for CD11b enrichment or proceeded directly to antibody labeling and analysis. For CD11b enrichment, all cells were resuspended in 2mLs FACS buffer and 45μLs of CD11b Mouse Magnetic Microbeads (Miltenyi) were added before incubating for 20mins. Samples were diluted with 2mLS FACS buffer before following steps to enrich CD11b$^+$ cells using MS Columns (Miltenyi, 130-042-201). It is important to note that cell suspension should be added to the column immediately following initial rinsing with 500μL of FACS buffer to prevent clogging. CD11b enriched or unenriched cells were resuspended in 100μL FACS buffer with ZombieUV (1:100) prior to adding anti-CD45 FITC, anti-CD11b AF647, anti-Ly6G BV785, and anti-TER119 PE according to Supplemental Table 8. Samples were washed with 1mLS FACS buffer and then resuspended in 300μL FACS buffer. Sorting of TER119$^+$ and TER119$^-$ macrophages was done on a BD Fusion 2 into FACS tubes coated with FACS buffer. Following sort, cells were centrifuged and resuspended in cold DPBS for transfer to protein lobind Eppendorf tubes. Cells were counted and then 2 biological replicates were combined into one to yield enough protein for mass spectrometry analysis. This resulted in input of 4 WT mice and 8 JAK2 KO mice, yielding one biological replicate of WT TER119- and TER119+, and four biological replicates of JAK2 KO TER119- and TER119+ macrophages. Samples were stored at -80°C prior to being processed for MS following protocol for WP MS using the Preomics iST96x kit.

**Preparation of single cell suspension from 4T1 tumors**

Extracted tumors were manually broken down using a razor blade. Tumors were then transferred to 50mL falcon tube using RPMI (Cytiva). Samples were centrifuged at 500xg for 5mins, and supernatant discarded prior to addition of 12mLs of digest buffer consisting of RPMI, 10ug/mL DNase (Roche, 10104159001) and 25ug/mL Liberase (Roche, 5401054001). Tissue was incubated with buffer for 15mins at 37°C, rotating, then passed through a 40μm Cell Strainer cell strainer into a falcon tube. Digest and filtering were repeated to yield single cell suspension. Pellet was resuspended in 2mLs of ACK Lysing Buffer (Gibco, #A10492-01) and incubated 2mins at RT. 18mLs of RPMI with 10% FBS was added, and samples centrifuged 500xg for 5mins before aspirating supernatant. Samples were then washed 1x with FACs buffer prior to labeling with anti-CD11b APC, anti-CD45 FITC, anti-Ly6G BV785, anti-EpCAM PE, and ZombieUV (1:100) in 100μL of FACS buffer on ice for 30mins. Details of antibody dilutions can be found in Supplemental Table 8. Following labeling, samples were analyzed and sorted on a BD Fusion 2, isolating macrophages that were EpCAM$^-$ or EpCAM$^+$ into FACS buffer coated FACS tubes. All samples were washed with PBS following sorting and then snap frozen and stored at -80°C. Samples were prepared for MS analysis following protocol for WP MS using the Preomics iST96x kit.

**Mass Spectrometry Sample Preparation – Whole Protein**

Snap frozen cell pellets of sorted cells were allowed to thaw on ice before preparation with PreOmics iST96 kit. PreOmics iST lysis buffer in combination with boiling solubilizes membrane and soluble proteins.

**Mass Spectrometry Sample Preparation – Surface Enrichment**

Following FACS sorting and DPBS washing, cells were then washed 3x with DPBS at pH 8.0 and resuspended at a concentration of $25x10^6$/mL in DPBS pH 8.0. EZ-Link™ Sulfo-NHS-LC-Biotin (Thermo Fisher Scientific, A39257) was added to a final concentration of 1.67mM before incubation for 30mins at RT. The reaction was quenched by washing 3x with DPBS pH 8.0 supplemented with 100mM glycine. Samples were then pelleted and snap frozen for storage at -80°C.

For enrichment, samples were allowed to thaw briefly on ice and then 50μL of Preomics Lyse buffer was added before heating at 95°C for 10mins with shaking. A BCA assay was used to determine protein content and samples were normalized across replicates (Pierce, Thermo Fisher Scientific, 23225). Protein was precipitated out of lysis buffer with the addition of 2x volume ice-cold methanol, 0.5x volume ice-cold chloroform, and 1x volume of water. Samples were vortexed after the addition of each component followed by a 10min centrifugation at max speed (>20,000rcf) leading to a phase separation with the white protein disk in the middle. The bottom layer was removed using a gel-loading pipette tip (VWR, 76321-564) and top layer decanted allowing the protein disk to adhere to the side of the Eppendorf tube. Sample was then resuspended in 2x the original volume of methanol and bath sonicated for 10mins to solubilize. Samples were stored overnight at -80°C.

Samples were allowed to come to RT and then centrifuged at max speed (>20,000rcf) for 10mins. Methanol was removed and samples were allowed to air dry for 15mins. The following volumes assume an input of 500ug protein and were adjusted according to normalized concentrations for each sample. Protein was re-solubilized in 250μL freshly prepared 6M Urea in DPBS with 10μL 10% SDS in PBS (w/v) and sonicated for 30 secs, 1 sec on/1 sec off. An additional 60μL of 10% SDS in PBS was added and the sample was vortexed to incorporate. Samples were transferred to falcon tubes and diluted with 3mLs of PBS before adding 70μL of washed streptavidin beads (Thermo Fisher, PI29202). Samples were placed on a rotator and allowed to incubate for 2hrs at RT. Beads were pelleted by centrifuging for 2mins at 1600rcf and washed with 5mLs each of the following reagents, centrifuging and aspirating between each: 0.2% SDS in DPBS, 1M NaCl in DPBS, 50mM ammonium bicarbonate (ABC) in 2M Urea. 500μL of DPBS was used to transfer samples to a new Eppendorf tube. An additional 500μL was used to wash the tube and transferred to the same Eppendorf before repeating spin and aspiration. Pelleted beads were resuspended in 70μL of Preomics LYSE and 50μL of Preomics DIGEST and incubated for 2hrs at 37°C with shaking. Beads were transferred to bio-spin columns (Bio-Rad, 7326204) and centrifuged 1min at 800rcf. Resulting flowthrough was quenched with 120μL Preomics STOP buffer before transferring to a Preomics CARTRIDGE and following their standard purification protocol to yield vacuum dried peptides. Peptides were resuspended in Preomics LC-Load and run on a Bruker timsTOF Pro.

**Mass Spectrometry Data Acquisition and Search Parameters**

All raw mass spectrometry data files are available upon request.

A nanoElute was attached in line to a timsTOF Pro equipped with a CaptiveSpray Source (Bruker, Hamburg, Germany). Chromatography was conducted at 40°C through a 25cm reversed-phase C18 column (PepSep, 1893476) at a constant flowrate of 0.5μL $min^{-1}$. Mobile phase A was 98/2/0.1% water/MeCN/formic acid (v/v/v, Thermo Fisher, LS118, LS120) and phase B was MeCN with 0.1% formic acid (v/v). During a 108 min method, peptides were separated by a 3-step linear gradient (5% to 30% B over 90 min, 30% to 35% B over 10 min, 35% to 95% B over 4min) followed by a 4min isocratic flush at 95% before washing and a return to low organic conditions. Experiments were run as data-dependent acquisitions with ion mobility activated in PASEF mode. MS and MS/MS spectra were collected with *m/z* 100 to 1700 and ions with *z* = +1 were excluded.

Raw data files were searched using PEAKS Online Xpro 1.7.2022-08-03_160501 (Bioinformatics Solutions Inc., Waterloo, Ontario, Canada). The precursor mass error tolerance and fragment mass error tolerance were set to 20 ppm and 0.03 respectively. The trypsin digest mode was set to semi-specific and missed cleavages was set to 2. For experiments with human cells, the human Swiss-Prot reviewed (canonical) database (downloaded from UniProt) and the common repository of adventitious proteins (cRAP, downloaded from The Global Proteome Machine Organization) totaling 20,487 entries were used. For mouse samples, the mouse Swiss-Prot reviewed (canonoical) database totaling 17,053 entries was used. Carbamidomethylation was selected as a fixed modification. Oxidation (M) and Deamidation (NQ) was selected as variable modifications. When relevant, data was searched with 13C(6)15N(2) and 13C(6)15N(4) SILAC labels as fixed or variable modifications.

**Mass Spectrometry Data Curation**

***4T1:*** Data was searched as described with a mouse FASTA. $EpCAM^+$ and $EpCAM^-$ macrophages datasets were normalized and ratios for each paired sample replicate were generated. The median ratio was generated (zeros were excluded) across 3 replicates with a maximum of 100 and a minimum of 0.01. To generate the dataset of proteins of putative 4T1 origin, we required a protein have a median area in $EpCAM^+CD45^-CD11b^-CD14^-Ly6G^-$ to be at least 500 and be at least 4-fold more detectable in this cell type compared to $CD45^+CD11b^+CD14^+Ly6G^+EpCAM^-$ macrophages. We further required that $EpCAM^+$ and $EpCAM^-$ enrichment occur in at least 2 of 3 replicates.

***Red Blood Cells:*** Proteins were required to be detectable in samples from 4 of 6 mice.

***Ter119-positive and Ter119-negative Spleen-Resident Macrophages***: Data was searched as described with a mouse FASTA. $Ter119^+$ and $Ter119^-$ macrophages datasets were normalized and ratios for each paired sample replicate were generated. The median ratio was calculated across all replicates with a set maximum of 100 and a set minimum of 0.01. To be considered enriched, we required the protein to be enriched in at least 3 of 8 replicates.

***Comparative Surface Enrichment***: For comparative surface enrichment analyses, data were searched using PeaksOnline as described with a surface-specific human FASTA. For isotopically 'light' surface datasets, area per protein was normalized with respect to total area per replicate. For isotopically 'heavy' surface datasets, area was not normalized. Median ratios were generated for Eaters and Non-Eaters across 4 technical replicates and a median ratio of Eaters/Non-Eaters was calculated. Proteins were denoted as 'Up in Eaters' if they were at least 4-fold more detectable in Eaters compared to Non-Eaters. Proteins were denoted as 'Up in Non-Eaters' if they were at least 4-fold more detectable in Non-Eaters compared to Eaters. For the 'heavy' surface protein dataset, proteins were cross-referenced with surface enrichment datasets from naive macrophages and SW620 cells.

**FACS Sorting and RNA extraction for RNA Sequencing**

Cell sorting was performed as described above (FACS Sorting for Mass Spectrometry Analysis). Following PBS washing steps to remove excess FBS, samples were lysed into TRI Reagent (Zymo Research, R2071) and then snap frozen and stored in -80°C until RNA extraction was performed. RNA extraction was performed using Direct-zol RNA MiniPrep Plus (Zymo Research, R2071) following manufacturer's instructions. In brief, an equivalent volume of 200 proof Ethanol was added to samples in Tri Reagent. The mixture was transferred to a Zymo-Spin IICR Column in a collection tube and centrifuged. In a separate tube, 5μL of DNase I was mixed with 75μL of DNA Digestion Buffer. 400μL of RNA Wash Buffer was added to the column and centrifuged prior to the addition of 80µL of DNaseI solution. Samples were incubated at RT for 15mins. Washing of the RNA with RNA Wash buffer was repeated 2 more times, first with 400µL and then with 700µL of buffer, centrifuging and removing buffer between each step. Columns were transferred to new RNase-free tubes and RNA eluted with 50µL of DNase/RNase free water. Samples were quantified via nanodrop and stored in -20C prior to sequencing.

**RNA Sequencing of Eaters, Non-Eaters, Uneaten, SW620 and Naïve Macrophages**
Two biological replicates of each cell type (Eaters, Non-eaters, Uneaten, SW620, and Naïve macrophages) were used for RNA sequencing, with each biological replicate consisting of two technical replicates. RNA-seq libraries were prepared using the Single Cell/Low Input RNA Library Prep Kit (NEB, E6420), following the manufacturer's protocol. In brief, 200ng of total RNA was used as input. First, reverse transcription was performed, followed by 7 cycles of cDNA amplification and clean-up. The cDNA was then fragmented, repaired, ligated with NEBNext adaptors, and enriched using 4-5 cycles of PCR. Libraries were sequenced on a NovaSeq X 10B platform using paired-end 150 bp (PE150) reads.

**RNA Sequencing Data Curation**
RNA-seq reads were trimmed to remove sequencing adapters using Trimmomatic (v0.39) with the TruSeq3-PE-2 adapter sequences. Low-quality bases and short reads were filtered using the parameters LEADING:3, TRAILING:3, SLIDINGWINDOW:4:15, and MINLEN:36. Trimmed reads were aligned to the human reference genome (hg38) using STAR (v2.7.11b). Gene expression levels were quantified using HTSeq-count, where aligned reads were counted against gene features defined in the Gencode v45 GTF annotation. Gene expression counts were processed with the PyDESeq2 library for differential expression analysis. Genes with total read counts below 10 were filtered out to reduce noise and focus on biologically relevant genes. The DESeq2 pipeline was employed to normalize the raw count data and account for differences in sequencing depth. Log-transformed normalized counts were calculated using the natural log of the normalized counts plus one (log1p transformation), ensuring that no zero values were present.

**Preparation of Primary Human Monocyte-Derived Macrophages**
Primary human donor-derived macrophages were generated as described previously[3,4]. Patient blood was received from anonymous donors in Leukocyte Reduction System Chambers (LRSC) from Vitalant Blood Donation. All reagents and materials which contact blood are sterilized with 10% Bleach solution prior to disposal in biohazard waste. Monocytes are isolated through sequential density gradients as described below. Critical equipment is a pipet aid capable of slow/gravity drain and a centrifuge where the brake can be fully disengaged.

The bottom of the LRSC was cut and entire chamber set into a 50mL polypropylene Falcon Tube prior to cutting the top to allow for gravity drain of the blood (anticipate 5-10mLs). Sample was diluted up to 35mLs with autoMACS Running Buffer (Miltenyi Biotec, 130-091-221) and gently rocked until fully mixed. The first density gradient was set using a serological and pipet aid (Drummond, SLOW dispense) to add 14mLs of Ficoll-Paque PLUS (Cytiva, GE17-1440-02) to the bottom of the tube. Sample was transferred to centrifuge (Beckman Coulter Allegra X-30) and balanced with falcon tube containing water which provided better separation compared to balancing two falcon tubes containing blood. Sample was spun at 400xg for 42mins with acceleration set at 4 and brake set at 0.

Post-centrifuge, sample contained 4 independent layers from top to bottom: excess autoMacs and serum, buffy coat, excess ficoll-paque, and pelleted red blood cells. The sample was carefully transferred to the tissue culture hood and the top serum layer was gently aspirated without disrupting the buffy coat. Serological pipette was used to remove white buffy coat layer and transfer to new 50mL falcon tube prior to dilution up to 50mLs with autoMACS. The sample was centrifuged at 400xg for 10mins with full acceleration and brake. Supernatant was aspirated and pellet was resuspended in 1mL of autoMACs buffer using a 1mL pipette before repeating the dilution with autoMACs and centrifugation. After aspirating, pellet was resuspended with 25mLs of RPMI Complete. Pellet was resuspended with 1mL by a pipette prior to addition of the remaining 24mLs using a serological. Percoll mix was prepared for second density gradient by mixing 3.47% HBSS 10x (Corning, 21-022), 42.55% Percoll (Cytiva, **17089102**), and 53.98% RPMI Complete (15mLs required per LRSC). Serological and pipet aid (SLOW dispense) was used to add 15mLs of Percoll Mix to the bottom of the falcon tube containing diluted sample prior to centrifugation at 400xg, 42mins, 4 acceleration, and 0 brake.

Post-centrifuge, the sample again contains 4 independent layers with the white center buffy coat being the isolated monocytes. The top layer is carefully aspirated and serological is used to collect the delicate disk of cells. Washes described above (up to 50mLs autoMacs, 400xg, 10mins) were repeated. Pellet was resuspended in 1mL autoMacs and 10µL was removed for count. The 10µL of cells was sequentially diluted (1:100, 1:10) for a final dilution of 1:1000. 10µLs of the final dilution was mixed 1:1 with Trypan Blue Stain (0.4%, Invitrogen, T10282) and cells counted via hematocytometer (recommend manual counting due to impurity). Sample will likely contain a mixed population of larger cells (monocytes) and very small cells (platelet or RBC contaminants). Count should only include larger cells. Sample should be spun and autoMacs aspirated prior to resuspension in Hyclone IMDM (Cytiva, 16777-370) supplemented with 2mM GlutaMAX at 20-30 million cells/mL. Standard Petri Dishes (Fisher, 89230-472) should be prepared with 10mLs of IMDM+GlutaMAX prior to the addition of 1mL of diluted monocytes. Sample should be incubated at 37°C, 5% CO2 for 45mins-1hr prior to aspirating media and replacing with 10mLs of IMDM+GlutaMAX with 10% Human Serum AB (Gemini Bio Products, 100-512-100). Day of plating is considered Day 0. Media exchanges should be performed on Day 3 and Day 5 by aspirating media, washing 3x with 5mLs of Dulbecco's Phosphate Buffered Saline, and replacing with 10mLs of IMDM+GlutaMAX+Human Serum. Interferon gamma stimulation was performed as described previously. In brief, 100 U $mL^{-1}$ of interferon-gamma was supplemented into IMDM+GlutaMAX+Human Serum on Day 5 and allowed to incubate at 37°C for 48-72hrs.

**Biotin-Labeling for Biotin Transfer Assay**

Following staining of immortalized cells of interest with CalceinAM as stated above, samples were counted via hemacytometer before being washed 3x with DPBS at pH 8.0. Samples were resuspended at a concentration of $25x10^6$/mL in DPBS pH 8.0. EZ-Link™ Sulfo-NHS-LC-Biotin was added to a final concentration of 1.67mM before incubation for 30mins at RT. The reaction was quenched by washing 3x with DPBS pH 8.0 supplemented with 100mM glycine. Samples were then washed 1x with DPBS pH 7.4 before being combined and co-incubated as stated above in "**Flow Cytometry-Based Phagocytosis Assay**". Following co-incubation, cells were washed 2x with FACS buffer and then resuspended in 100µL FACS buffer with anti-CD11b AF647 and Streptavidin BV711 according to Supplemental Table 8. Samples were incubated on ice for 30mins before washing with 1mL FACS buffer, centrifuging, and resuspending in FACS buffer with Sytox Orange (1:10,000) viability dye and incubated 10mins at RT prior to analysis on Cytek Aurora Spectral Flow Cytometer.

**SLC1A5 Labeleing and Homopropargylglycine (HPG) Uptake Assay**

HPG update assay was modified from Pelgrom *et al*[5]. CellTracker Green CMFDA (Invitrogen, C7025) was allowed to come to RT. Staining working stock was produced by adding 10.8µL of fresh DMSO to 20ug aliquot of CellTracker Green (10mM). Stock was diluted into IMDM (Cytiva) supplemented with 2mM Glutamax (Gibco) but with no serum added for a final concentration of 1µM. Media was removed from SW620s which were plated two days in advance and allowed to reach 80-90% confluency. Plates were washed 1x with DPBS and then 10mLs of IMDM with CellTracker Green (1µM) was added and incubated at

37°C, 5% CO2 for 20mins. IMDM was removed and plates were washed 2x with FBS containing media (SW620 culture media listed above) and 2x with DPBS prior to lifting with TrypLE prior to counting. Stained SW620s were co-incubated with human monocyte derived macrophages as outlined in "**Flow Cytometry-Based Phagocytosis Assay**". Following co-incubation, cells were washed 2x with DPBS prior to labeling with Zombie Yellow Fixable Viability Kit (1:1000) in PBS. Cells were washed with FACS buffer prior to washing with DPBS.

In a 96-Well Clear Ultra Low Attachment U-Bottom Microplate, 80µL of 200µM HPG (Vector, CCT-1067) or 200µM HPG with 4mM glutamine (Fisher, NC1321615) was added and solutions were allowed to equilibrate to either 37°C or 4°C for 15mins. Following pre-incubation, combined SW620s and macrophages (400,000:200,000) were aliquoted in PBS in 50µL into same 96-well plate. Multichannel was used to add 50µL of equilibrated HPG or HPG with glutamine to the 50µL of cells. Plates were then incubated for 5mins at 37°C or 4°C as indicated within figures. Reaction was quenched by placing all plates on ice and then fixing by incubating 15mins with 100µL of 2% PFA (Biotium, 22023) at RT. Samples were washed 1x with DPBS and then 200µL of 0.01% Digitonin (Sigma, D141) in PBS was added and samples were incubated 5-10mins at RT to permeabilize plasma membrane but retain internal membrane structures[6]. During fix and permeabilization, the click chemistry mixture was prepared with 1 part CuSO4 (Sigma, C7631, 100mM in MilliQ), 1 part NaAsc (Sigma, A4034, 1M in MilliQ), 1 part THPTA (Vector, CCT-1010, 100mM in MilliQ), 96 parts DPBS (Cytiva), and 0.25 parts of Az-AF594 (Vector, CCT-1295, 2mM in DMSO). Samples were washed 2x with PBS and then incubated for 1hr at RT in the dark with 30µL of prepared click mixture. Following incubation, samples were washed with FACS buffer and then incubated for 30mins, RT, in the dark with 200µL of 55mM EDTA (Fisher, S25311A, 55mM in PBS). Samples were washed a resuspended in 100µL FACS buffer for staining with anti-SLC1A5 antibody for 30mins on ice. Following washing, final staining was performed in 100µL FACS with anti-rat IgG2a BV421 and anti-CD11b AF647. All antibodies are added according to Supplemental Table 8. Samples were washed and resuspended in FACS buffer for analysis on Cytek Aurora.

**Preparation of Single Cell Suspension of Primary Human Tumors**

Collection of human tumor tissue was approved by UCSF IRB and patients de-identified prior to experimental use. Samples were stored in PBS at RT until beginning digestion. Tissue was transferred to C tubes (Miltenyi) with 2-3mLs of digestion buffer and scissors were used to manually break down the sample. C tubes were placed into gentleMACS (Miltenyi) for 1hr at 37°C to fully dissociate the tissue and yield a single cell suspension. Dissociated cells were washed with FACS buffer and then hemocytometer was used to measure cell concentration.

Donor matched PBMCs were isolated from patient blood. Collected blood was transferred from collection tube into a 15mL Falcon Tube and diluted up to 13mLs with ACK Lyse Buffer. Samples were incubated for 7mins at RT before centrifugation and removal of supernatant using a serological. ACK Lyse treatment was repeated prior to washing cells with FACS buffer.

Cells were stained with a fixable viability dye (Zombie Yellow, Ghost Dye Red 780) and either fixed with 2% PFA in PBS for 15mins at RT or proceeded directly to labeling with anti-CD11b AF647, anti-CD14 FITC, anti-EpCAM PE for 30mins on ice according to Supplemental Table 8. Fixed samples were stored in 4°C overnight and then labeled with surface antibodies and analyzed on Cytek Aurora the following day.
